# Exhaustive Cross-linking Search with Protein Feedback

**DOI:** 10.1101/2022.03.03.482813

**Authors:** Chen Zhou, Shuaijian Dai, Yuanqiao Lin, Ning Li, Weichuan Yu

**Affiliations:** Department of Electronic and Computer Engineering, The Hong Kong University of Science and Technology, Hong Kong; Division of Life Science, The Hong Kong University of Science and Technology, Hong Kong

**Author notes:** Authors contributed equally to this paper (Chen Zhou, Shuaijian Dai, Yuanqiao Lin).

## Abstract

Improving the sensitivity of protein-protein interaction detection and protein structure probing is a principal challenge in cross-linking mass spectrometry (XL-MS) data analysis. In this paper, we propose an exhaustive cross-linking search method with protein feedback (ECL-PF) for cleavable XL-MS data analysis. ECL-PF adopts an optimized *α/β* mass detection scheme and establishes protein-peptide association during the identification of cross-linked peptides. Existing major scoring functions can all benefit from the ECL-PF workflow to a great extent. In comparisons using synthetic datasets and hybrid simulated datasets, ECL-PF achieved three-fold higher sensitivity over standard techniques. In experiments using real datasets, it also identified 91.6% more cross-link spectrum matches and 52.6% more unique cross-links.

## Introduction

Cross-linking mass spectrometry (XL-MS) is used to deduce protein-protein interactions (PPIs) and decipher protein structures in a high-throughput manner [1–4]. In the XL-MS technique, cross-linkers are designed to produce spatial information in a cellular context. A cross-linker consists of two reaction groups connected by a spacer-arm. In experiments, both reaction groups link amino acid residues to form covalent bonds under the constraint of the spacer arm’s length. Mass spectrometry (MS) is then used to identify the cross-linked residues so as to provide information about PPIs and protein structures. Cross-linkers can be divided into non-cleavable cross-linkers [5–7] and cleavable cross-linkers [8–11]. Non-cleavable cross-linkers involve “*n*-square” time complexity during the identification of cross-linked peptides; that is, the number of possible peptide pairs satisfying the precursor mass constraint is quadratic to the number of linear peptides in the database. To reduce the quadratic time complexity, cleavable cross-linkers were invented. A cleavable cross-linker breaks into two residues in the collision-induced dissociation (CID) process and generates signature ions in the MS/MS (MS2) spectrum, which can help us directly pinpoint the *α/β* peptide mass. Although state-of-the-art software programs are capable of analyzing non-cleavable cross-linking data in linear time complexity [12, 13], the speed is still 10^3^ ∼ 10^5^ times slower than that of the cleavable cross-linking task (Supplementary Section 1). In this paper, we will thus focus on the analysis of cleavable cross-linking data.

Though the XL-MS approach seems straightforward, identifying cross-linked peptides is challenging. The major obstacle is the unequal fragmentation efficiency of the cross-linked peptide pair. This imbalanced fragmentation [14, 15] in the CID process always makes one of the cross-linked peptides fragment into fewer ions or even no ion, resulting in many of the MS2 spectra being unidentifiable and thus hampering sensitivity. Different strategies have been proposed to generate more useful ions, such as utilizing an electron-transfer dissociation (ETD) MS2 spectrum [16], stepped higher-energy collision dissociation (stepped-HCD) MS2 spectrum [17], or CID MS/MS/MS (MS3) spectrum [18, 19]. These approaches try to take advantage of the hardware. From the perspective of software development, however, only a handful of work has been done so far [16, 20–24], and the sensitivity is still in the range of 20% ∼ 50% (observed from simulated datasets). We have discovered that using protein-peptide association during the identification process improves the sensitivity considerably. In addition, we demonstrate that major scoring functions in XL-MS can all benefit from this protein-peptide association with a large margin.

Here, we describe a flowchart that exhaustively searches all possible cross-linked peptide pairs and propose a novel protein feedback pipeline that discovers the correspondence between protein and peptide to improve the sensitivity of the search engine. We demonstrate the generalizability and effectiveness of this feedback mechanism with several major scoring functions. Adopting the ECL-PF design, we have observed significant sensitivity improvements from different types of datasets, including synthetic datasets, hybrid simulated datasets, and *in vivo/vitro* real experimental datasets. The result shows that ECL-PF can uncover more intriguing information on PPIs and protein structure at no cost of precision.

## Methods

### Exhaustive cross-linking search

Cleavable cross-linkers are designed to be fragmented during the dissociation process in order to provide signature ions to help indicate the mass region of the cross-linked *α* and *β* peptides. Here, we define the peptide with the heavier mass in the cross-linked peptide pair as the *α* peptide and the lighter one as the *β* peptide. Signature ions should be robustly detected to decipher the individual peptide mass. Otherwise, even a good-quality spectrum can be unidentifiable and wasted. We summarize current approaches to determining the *α/β* peptide mass and describe a flowchart that exhaustively searches every possible candidate to pinpoint the peptide mass under the assumption that at least one signature ion should should up. More than that, a local alignment module is designed for compensation when no signature ion is available in the current MS2 spectrum. According to our comparison, this compensation can provide ∼ 10% more cross-link spectrum matches (CSMs) (Supplementary Section 2).

### Protein feedback pipeline

Due to unequal fragmentation in XL-MS, algorithms often need to identify poor fragmented peptides. Designing suitable scoring functions for this situation can help find correct peptides but won’t solve the problem entirely, especially under the condition that no fragmented ions of the *β* peptide are available. Scoring functions can not robustly identify low-quality spectra alone in XL-MS. Thus, we consider using more information, namely, protein feedback, to help the scoring function identify the low-quality spectra.

In the following, we provide a detailed description of the protein feedback pipeline of ECL-PF, which is shown in Figure 1(a). In the first step, the query spectra are directly matched by the scoring function *S*(*x*). The scoring function stores the top *p* peptide candidates and their corresponding scores for further judgment. In ECL-PF, we provide three major alternative scoring functions for users to choose: a simplified XlinkX scoring function [16, 20] given in Eq. 1, a normalized MeroX scoring function [21, 22] given in Eq. 2, and an Xcorr scoring function (SEQUEST) [26] given in Eq. 3:

**Figure 1:**
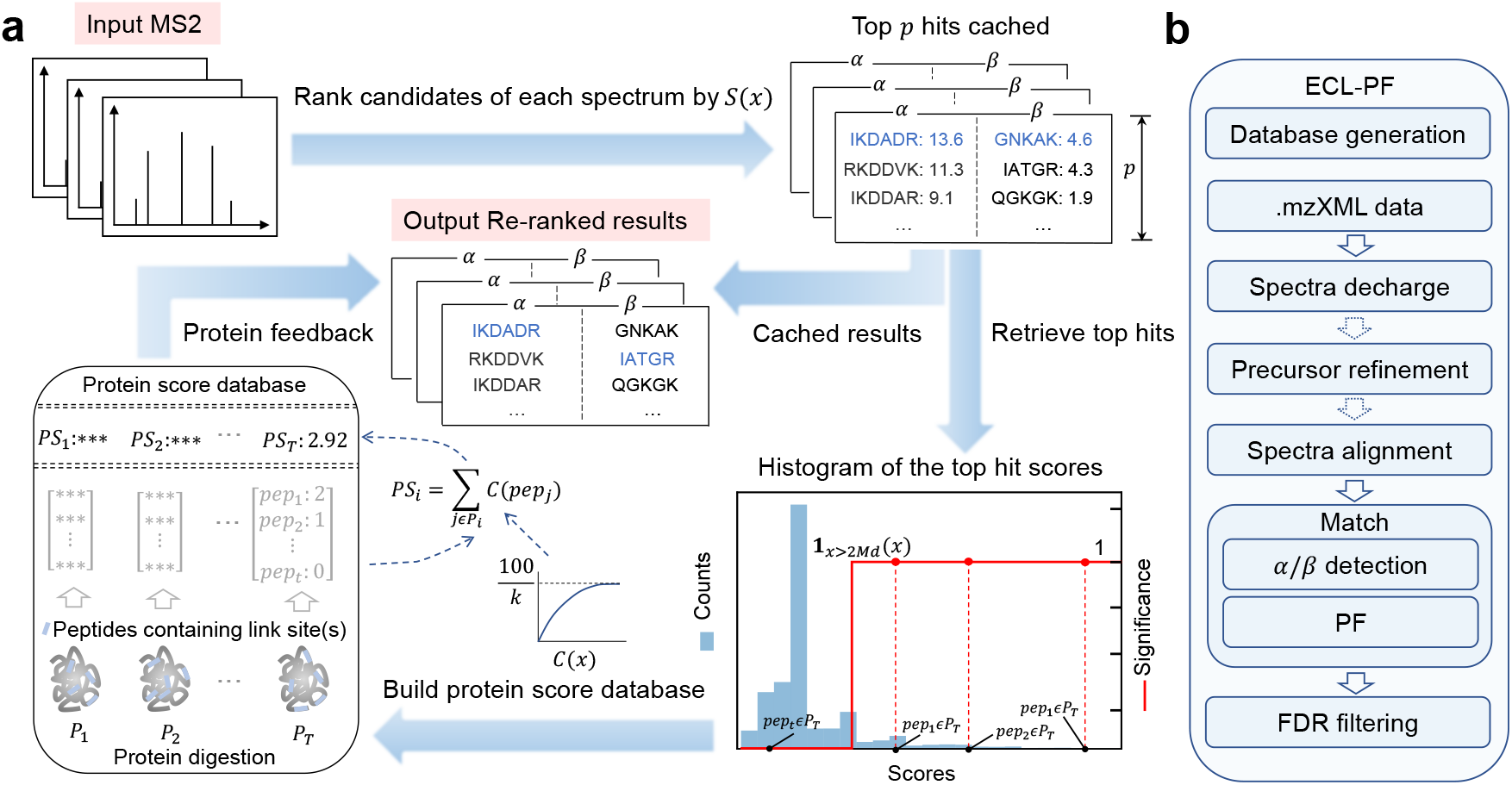
Protein feedback pipeline and general workflow of ECL-PF. (a) Details of protein feedback pipeline. The input MS2 spectra are first matched with peptide sequences by the scoring function *S*(*x*). The algorithm stores the top *p* hits for each spectrum during the matching process. We then retrieve the first *α* and *β* hits of all the spectra and plot the histogram of the score distribution in a further step. Adopting an indicator function (2*Md* is twice the median number), we build a protein score database using the filtered (significant) peptides. Each protein in the database is first digested into peptides, and we only use the peptides containing a link site(s) to construct a vector for that protein. Next, we use the counting function *C*(*x*) to compute the protein score. In the last step, we go back to the stored top *p* peptides and find the largest protein score among these candidates so as to finish the matching process and output the results. (b) General workflow of ECL-PF. Peptide database generation is separated from the spectra matching. The data are first deisotoped by Hardklör [25]. Then running a precursor mass refinement function is suggested to recalculate the precursor mass for each spectrum. Spectra alignment is also recommended when dealing with a small/purified dataset. A further step is the *α/β* peptide detection scheme and protein feedback pipeline. Last, ECL-PF conventionally utilizes target-decoy FDR filtering to export the cross-linked result.

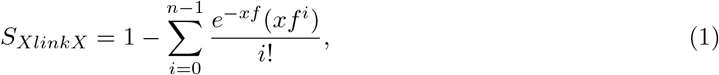

here *x* = 1/111.1 × 2 × (*M*_*prec*_ × *ϵ* × 2); *ϵ* is MS2 tolerance (ppm); and *f* is equal to *α/*(*α* + *β*) × *f*_*total*_ when we calculate *α* peptide score or is equal to *β/*(*α* + *β*) × *f*_*total*_ when we calculate *β* peptide score, where *α* and *β* denote *α* and *β* peptide mass and *f*_*total*_ is the total number of fragmented ions in the MS2 spectrum.

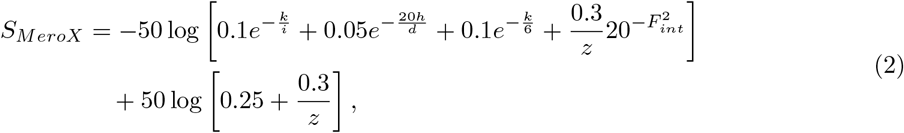

here *k* is the number of identified abundant ions (≥ 10% intensity); *i* is the number of abundant peaks in the MS2 spectrum; *h* is the number of identified ions; *d* is the total fragmented peaks in the MS2 spectrum; and *F*_*int*_ is the ratio of the intensity of identified ions to that of total ions in the MS2 spectrum.

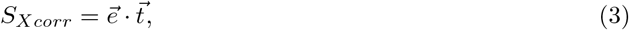

here 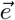 is the vectorization form of the experimental MS2 spectrum and 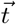 is the vectorized theoretical peptide sequence generated from the *b/y* ions.

After scoring peptides, the algorithm retrieves the top *α* and *β* hit of each spectrum and draws the histogram of their scores in the second step. A well-designed scoring function should have the property that the higher the scores, the more confident the peptides. We employ an indicator function to group the peptides by their scores in a binary way. Empirically, the significant domain is twice the median or 98th percentile of the scores. All the significant peptides are then used to build a protein score database.

We introduce a novel concept of protein score database as follows. Each protein in the FASTA file is *in sillco* digested into peptides. We use the peptides containing link site(s) to build a vector for that individual protein. The vector length is the same as the peptide number. If one specific peptide of this protein is discovered in the significant area in the histogram *m* times, the corresponding position in the vector is *m*. For example, in figure 1(a), protein *T* can be digested into *t* distinct peptide sequences. Its vector length is *t*. In the histogram, sequence *pep*_*1*_ is identified twice, and sequence *pep*_*2*_ is identified once in the significant area. We then initialize the vector of protein *T* as shown in the protein score database. After enumerating all proteins in the FASTA file, we design counting function *C*(*x*) in Eq. 4 to transfer the counts and sum up all the counting results from the protein vector to obtain the final protein score *PS*. If protein *T* has 20 link sites in its sequence, the final score can be calculated as 2.92. We also use a concrete protein sequence to show how to calculate protein score in Figure S10. Different from peptide databases that are used to provide potential peptide sequences to identify experimental spectra, the protein score database is used to provide global information to connect peptides to proteins and is established by the theoretical protein sequence and the scoring function.

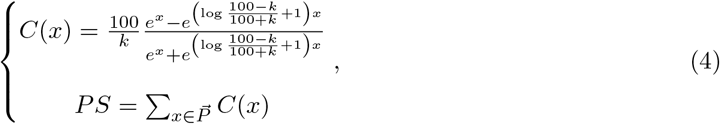

here *k* is the link site(s) number in the individual protein and 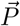 is the protein vector. The protein score *PS* is calculated by summing the counting result. The motivation of this function is explained in Supplementary Section 3.

The last step is the feedback procedure. The algorithm returns to the stored top hits and looks up the peptide candidates in the protein score database. Among the peptide candidates, we regard the one having the largest protein score as the peptide match for that spectrum. Testing on a proteome-wide dataset [27], we have found that the protein feedback pipeline can universally improve the performance of the scoring functions (*S*_*XlinkX*_, *S*_*MeroX*_, and *S*_*Xcorr*_) by 29.7%. More details of the protein feedback are elaborated in Supplementary Section 3.

### General workflow of ECL-PF

Currently, ECL-PF only handles cleavable cross-linking from CID(HCD)-MS2 spectra or CID(HCD)-MS2-ETD-MS2 spectra. It generates the peptide database separately from the matching process in the workflow (shown in Figure 1(b)). ECL-PF requires data in mzXML format as the input. Typically, we use the ProteoWizard tool (MSConvert) [28] to convert RAW data into mzXML format enabling peak picking, zero sample, and 32-bit binary encoding precision functions. ECL-PF then adopts Hardklör [25] to deisotope the MS2 spectra. It integrates Hardklör into its pipeline and provides a default parameter setting based on a high-resolution mass spectrometer. Users can then refine the precursor mass by running the mass refinement module in the ECL-PF. According to our experiments, this can improve the performance by ∼ 30% (Supplementary Section 4). A local alignment module is also recommended in small/purified datasets. After that, ECL-PF will proceed to the matching step using the *α/β* mass detection scheme and protein feedback pipeline. Finally, it employs a target-decoy strategy to filter the significant results. ECL-PF separately calculates the false discovery rate (FDR) of inter-protein cross-linked results and intra-protein cross-linked results by Eq. 5 [12, 29]:

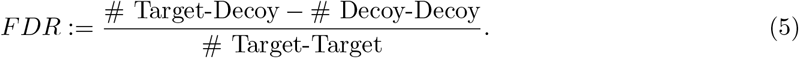

## Experimental results

### Evaluation using synthetic dataset

MS cleavable datasets were downloaded from the synthetic cross-linked peptide library (PXD014337) [30]. DSSO and DSBU were used to cross-link 95 synthesized peptides. PD 2.4 MS Annika, PD 2.4 XlinkX 60-day free version, MaxLynx (MaxQuant 2.0.3.0), and MeroX 2.0.1.4, as well as ECL-PF, are used to analyze these datasets. Local alignment and precursor mass refinement modules are enabled in ECL-PF. Table S8 shows the concrete settings.

The results for the number of CSMs and unique cross-links at FDR= 5% are shown in Figure 2 (FDR at 1% can be found in Figure S11). Among the CSMs results, ECL-PF always obtains the highest true positive numbers. On average, ECL-PF identifies 2.3 times as many CSMs as other software on the DSBU dataset and 3.0 times as many CSMs as the others on the DSSO dataset. For the unique cross-links, due to the limited number of synthesized peptides in the sample, ECL-PF does not outperform all the others but is in the same range as the best one. From the calculated FDR, we can observe that MS Annika suffers from a high proportion of false positives — on the DSSO dataset, one quarter of the unique cross-links are wrong. Therefore, we don’t compare it with the other software in the following sections. Meanwhile, it is observed that XlinkX obtains the worst results for all datasets. Thus, we also remove it from the comparisons hereafter.

**Figure 2:**
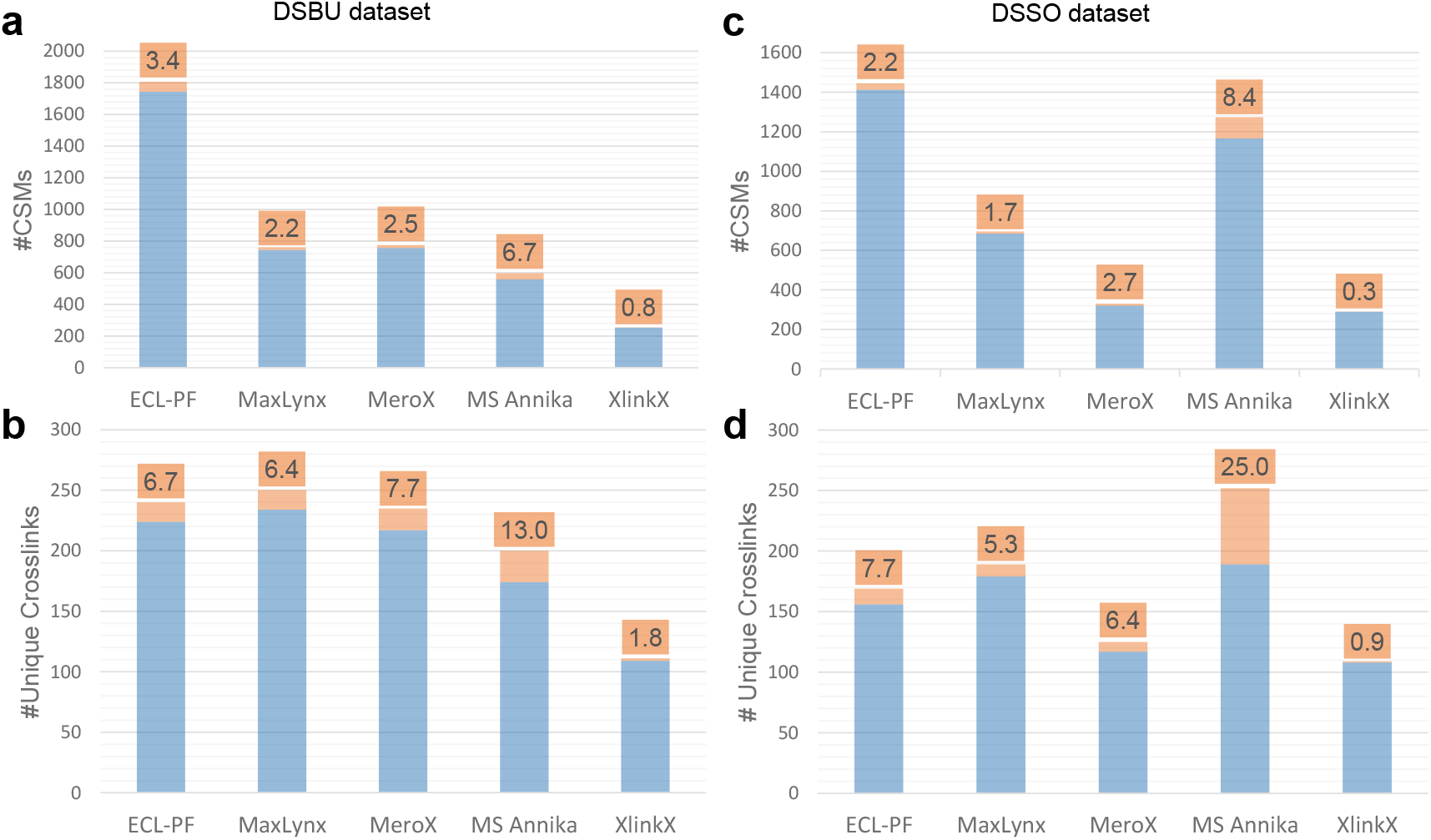
Software comparison using synthetic datasets. CSMs and unique cross-link results for DSBU (a)(b) and DSSO (c)(d) datasets are plotted. True positive results are denoted by a blue bar. False positives are denoted by an orange bar, and the proportion is at the top. On average, ECL-PF identifies 2 ∼ 3 times as many CSMs as other software. Due to the limited number of synthesized peptides in the samples, ECL-PF does not outperform all software programs in terms of the number of unique cross-links but is in the same range as the best one.

### Evaluation using hybrid simulated dataset

Due to the unique characteristics of XL-MS, we design a way to generate a simulated dataset based on a real experimental one. We separate the signal peaks of the *α* and *β* peptides from the noise peaks in each identified spectrum and then combine the *α* peaks from one spectrum with the *β* peaks from another spectrum to produce a new spectrum. In this way, if we originally have *N* identified spectra, we can generate *N* ^*2*^ simulated spectra with ground truth. Additionally, we can add their noise peaks into the new spectrum so as to have more realistic spectra. Here, we use the synthetic cross-linked peptide library as the source to generate the simulated dataset and tune the noise ratio to make the spectra more complicated. Supplementary Section 5 shows more details of the simulation.

MaxLynx cannot function with simulated datasets because its input only allows RAW files. Therefore, in this section, we compare ECL-PF only with MeroX. To fairly compare them, we use the overlapping spectra identified by both software to generate our simulated dataset. To test their robustness, we also add noise peaks into the spectra. In total, we generated 25,221 spectra for the DSSO cross-linker and 84,741 spectra for the DSBU cross-linker. Among them, we add the noise peaks extracted from the original spectra. The noise proportion varies from 0% to 100%, where 100% means we put all noise peaks from the original spectrum into the new spectrum. Figure 3 shows the sensitivity and precision comparison between ECL-PF and MeroX. ECL-PF identifies almost all the cross-linked spectra when there is no noise, whereas MeroX identifies half of them on average. As we increase the noise proportion, both of their sensitivities drop. ECL-PF can still identify more than 80% of the spectra in the worst case, but MeroX can only identify 20%. They both satisfy our FDR at the 5% setting. Simulated results show that ECL-PF can not only achieve better result but is also more robust to noise than MeroX.

**Figure 3:**
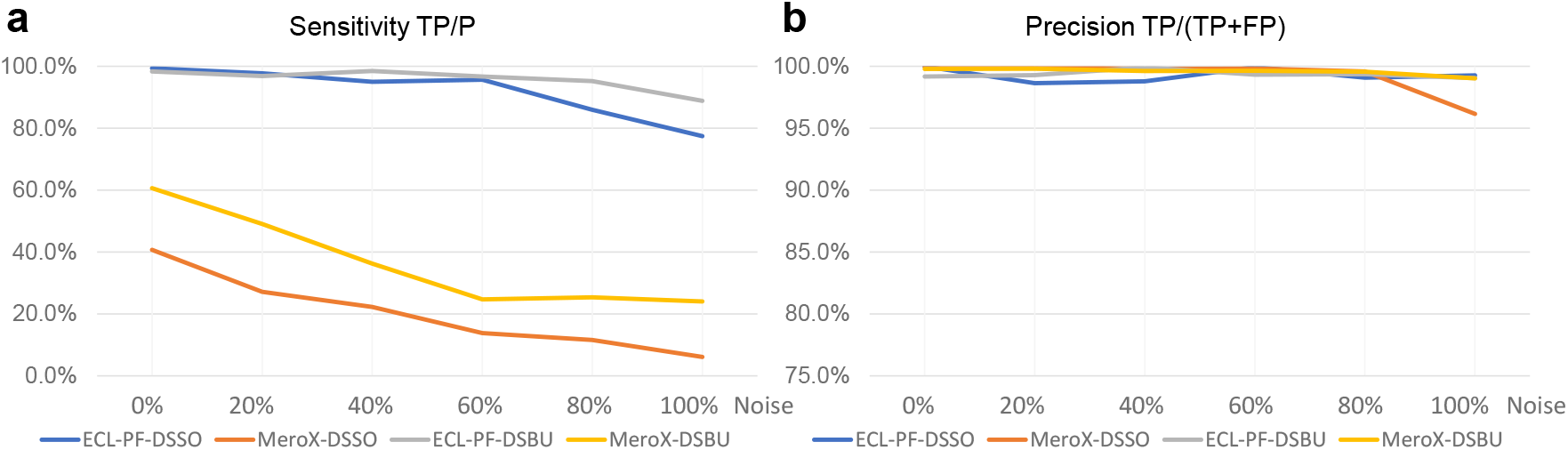
Software comparison using simulated datasets. The simulated dataset is derived from the DSSO and DSBU dataset. We add proportions of noise into the spectra, varying from 0% to 100% with a step of 20%. When there is no noise peak in the spectra, the sensitivity of ECL-PF is approaching 100%, whereas that of MeroX is about 50%. As more and more noise appears in the data, the sensitivity of both software drops. ECL-PF still identifies about 80% of the spectra in the worst case. In contrast, MeroX only identifies about 20% of the spectra for the DSBU data and less than 10% for the DSSO data. Most of the time, the precisions of both software are around 99%. When the noise level is increased to 100%, the precision of MeroX slightly drops to 96% for the DSSO data.

### Evaluation using real dataset

Four real datasets are used to evaluate the performance of ECL-PF, MaxLynx, and MeroX. The cross-linker varies from DSSO and DSBU to DSBSO. FDR at 1% is set for the four sets of comparisons. The precursor mass refinement module and local alignment module are enabled for ECL-PF. Supplementary Section 6 elaborates more information on the datasets and detailed settings. Table 1 shows the result of the comparisons. For all the tested data, ECL-PF outperforms the others by a large margin in terms of CSMs number and unique cross-links. On average, ECL-PF identifies 91.6% more CSMs and 52.6% more unique cross-links than the other methods in these real experimental datasets.

**Table 1:** Real dataset comparison among Maxlynx, MeroX and ECL-PF. PXD011861 [17], PXD016963 [31] and PXD020014 [32] are *in vitro* experimental datasets on an E.coli study. PXD012546 [27] is a proteome-wide *in vivo* dataset on a drosophila melanogaster study. The number of CSMs and unique cross-links are computed for each software on these datasets. At the bottom of the table, we calculate the average ratios of the software the to ECL-PF results.

| Data |  | MaxLynx |  | MeroX |  | ECL-PF |  |
| --- | --- | --- | --- | --- | --- | --- | --- |
|  |  | CSMs | Cross-links | CSMs | Cross-links | CSMs | Cross-links |
| *In vitro* experiments | PXD011861 | 4627 | 794 | 2511 | 672 | 5369 | 847 |
|  | PXD016963 | 1957 | 204 | 806 | 108 | 2665 | 333 |
|  | PXD020014 | 1888 | 101 | 613 | 78 | 4566 | 209 |
| *In vivo* experiment | PXD012546 | 31754 | 8211 | 28826 | 9422 | 48075 | 10260 |
| Ratio |  | 66.8% | 70.9% | 37.6% | 60.2% | 100% | 100% |

### Structure validation

E.coli GroEL complex results from ECL-PF were filtered from the real experimental dataset. We download the X-ray diffraction structure of the GroEL (PDB ID: 1KP8) and map our cross-linked results onto the protein structure. In total, 101 pairs of Lysine-Lysine sites are identified from ECL-PF, where 93.1% of the C*α*-C*α* distances satisfy the cross-linker’s constraint. The protein structure and histogram of the distances are shown in Figure 4. This structure study shows the solidity of ECL-PF from the experimental point of view.

**Figure 4:**
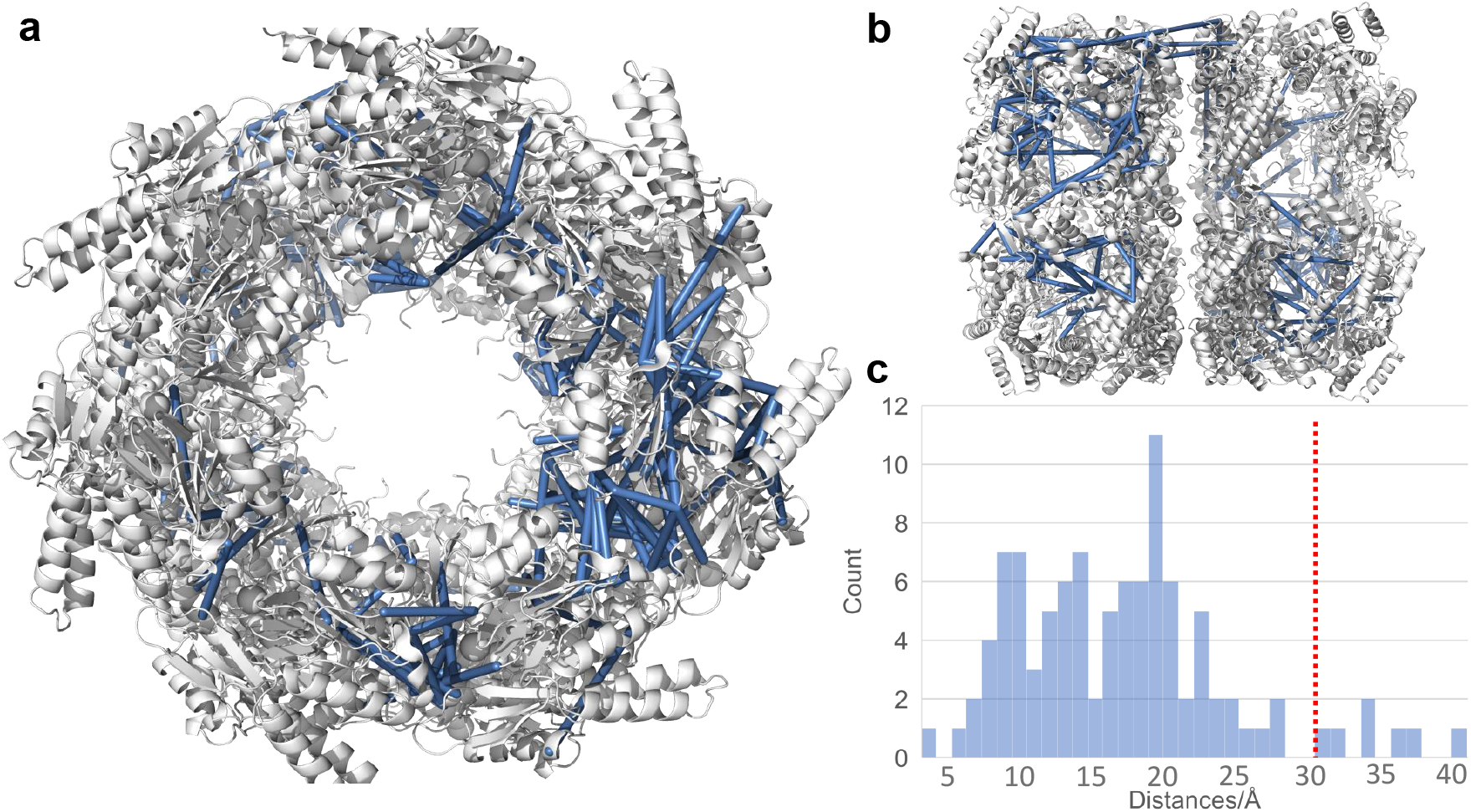
Protein structure validation. (a) The identified cross-links (solid blue lines) mapped onto the X-ray diffraction structure of the GroEL complex (PDB ID: 1KP8). (a) 90-degree rotation of the structure model in (a). (c) Histogram of *Cα*-*Cα* distance of all cross-links within the GroEL complex. 30Å is set as the distance cutoff shown in the red dotted line. 93.1% of the result satisfy the distance constraint.

## Conclusion

XL-MS presents powerful properties in PPIs and protein structure study. In this paper, we elaborated a comprehensive *α/β* peptide mass detection scheme and proposed a novel protein feedback pipeline to improve the sensitivity of current XL-MS software. Incorporating these ideas, we developed the ECL-PF program for cleavable XL-MS data analysis. Comparisons using synthetic datasets, hybrid simulated datasets, and real experimental datasets showed that the sensitivity of ECL-PF outperforms other software programs by a significant margin. Different scoring functions can also benefit from protein feedback. Potentially, protein feedback could be adopted in the non-cleavable cross-linking task as well.

## SUPPORTING INFORMATION

## 1 Time complexity analysis of non-cleavable cross-linking search and cleavable cross-linking search

We define the peptide with the heavier mass in the cross-linked pair as *α* peptide and the lighter one as *β* peptide. The mass of *α* peptide is denoted as *M*_*α*_ and that of *β* peptide mass is *M*_*β*_. Cross-linker mass and precursor mass are represented as *M*_*xl*_ and *M*_*prec*_, respectively.

Both non-cleavable search and cleavable search must satisfy Eq. S1.

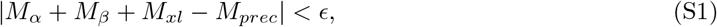

where *ϵ* is the mass tolerance. It is determined according to the mass spectrometer’s resolution.

To visualize Eq. S1, we *in silico* digest 100 proteins randomly chosen from the homo sapiens database into peptides by the trypsin rule, allowing two missed cleavages with a minimum peptide length of five amino acids and maximum mass of 6000 Dalton to draw a histogram according to the peptide mass in two-dimensional space, as shown in Figure S1. We treat the horizontal axis as the *α* peptide mass and the vertical axis as the *β* peptide mass. These two dimensions are asymmetric because we require *M*_*α*_ ≥ *M*_*β*_.

The grey area in Figure S1 denotes Eq. S1 and also indicates the naive exhaustive searching area for non-cleavable search. The red area means the cleavable cross-linking searching region. Benefitting from signature ions, the cleavable search engine can directly find a much smaller area of the target peptides. The zoom-in view on the right side of Figure S1 shows the searching area of the non-cleavable search engine (Xolik [1]) and cleavable search engine (XlinkX [2]) in linear time complexity.

To compare the time complexity of non-cleavable search with that of cleavable search in a numerical way, we neglect the data pre-processing procedure of the different software and assume the scoring function takes the same time to score every possible *α/β* peptides. In this way, we can directly use the area in the searching space to denote the time complexity of both the non-cleavable and cleavable cross-linking search methods by assuming the peptides are evenly distributed within the searching space.

We calculate the ratio of the grey area to red the area in Eq. S2:

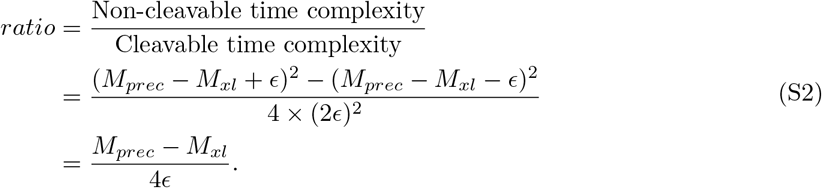

The ratio depends on several parameters. Precursor mass *M*_*prec*_ is often in the range of several thousand Dalton (∼ 10^3^), and *ϵ* is the mass tolerance depending on the device. Normally, *ϵ* ranges from 0.01 Da to 1 Da. Therefore, the ratio ranges from 10^3^ for large tolerance to 10^5^ for small tolerance.

The above numerical estimation motivates us to focus on the cleavable cross-linking search because it is more suitable for proteome-wide study in terms of speed.

**Figure S1:**
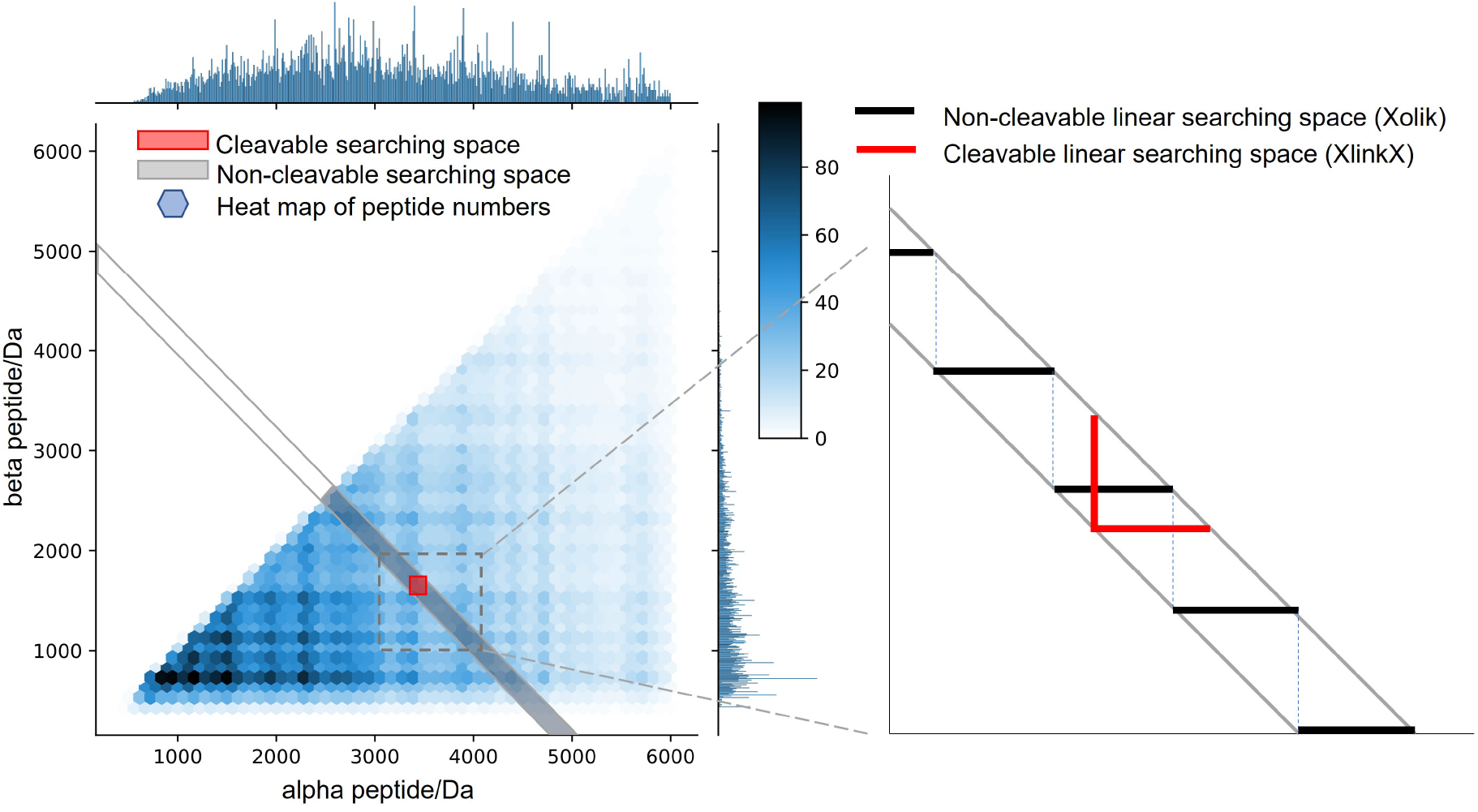
Visualization of searching space of non-cleavable cross-linking and cleavable cross-linking search. The heat map shows the joint distribution of the *α* peptide and *β* peptide. One hundred proteins randomly chosen from the human database are *in silico* digested by trypsin with two missed cleavage sites. The maximum mass is 6000 Dalton and the minimum peptide length is five. Only peptides containing Lysine are retained. The naive exhaustive searching method in non-cleavable cross-linking follows Eq. S1 with a further constraint that *M*_*α*_ ≥ *M*_*β*_, and is represented by the grey area. Cleavable searching methods ideally have the signature ions to target a specific location in the database, shown in the red area. We further draw the searching space of the non-cleavable search engine and cleavable search engine (Xolik and XlinkX) in linear time complexity on the right.

## 2 Exhaustive cross-linking search

Cleavable cross-linkers are designed to break during the dissociation process, providing signature ions to indicate the mass region of the cross-linked peptides. A naive way to use the signature ions is to calculate the mass difference between each pair of two peaks (known as a doublet) in the spectrum. If the mass difference is equal to the specific Dalton value, e.g., 32 Da for DSSO [3] and 26 Da for DSBU [4], we can treat the corresponding doublet as a pair of signature ions. Ideally, we can find two complementary doublets in a spectrum containing cross-linked peptides. Subtracted by the cross-linker’s residual mass, we can derive accurate masses for *α* and *β* peptides. However, due to the different settings of the mass spectrometer and different choices of the cross-linker, it is not always guaranteed that two complementary doublets will appear in the spectrum.

Some software considers the lack of complementary doublets and includes strategies to compensate for this situation. ReACT designs PIR [5] to generate a spike in the spectrum to indicate that the spectrum corresponds to cross-linked peptides. XlinkX is the first to propose delta mass strategy and top *N* intensity peaks’ strategy to derive the *α/β* mass.

We summarized all the situations in which the signature ions can appear and exhaustively search all the possibilities under the assumption that at least one peak should show up. Figure S2 shows every situation the signature ion(s) can appear in. We proceed with this flowchart and introduce each situation as follows:

1. For the query spectrum, we first pair up the peaks by the delta mass. Any two peaks will form a doublet if their distance satisfies the restriction according to the cross-linker. If more than one doublet exists, we compare any two of them to see if they are complementary to each other. This is the ideal situation that almost every algorithm will take into account.
2. The workflow will then proceed to check the second situation, in which the constraint is a doublet plus a singular peak equal to the precursor mass. In this situation, three signature ions are used to infer the *α/β* peptide mass.
3. In the third situation, we use just one doublet to derive the *α/β* peptide mass with the help of the cross-linker’s information and an accurate precursor mass.
4. After using all the doublets in the spectrum, we enumerate any two peaks to see if their summation is equal to the precursor mass plus/minus delta mass. Each pair of peaks that satisfies the constraint will proceed to situation four and the corresponding sub-situation.
5. Then we enumerate all the peaks again to see if any of the summations are equal to the precursor mass alone to see if they satisfy situation five. Note that the fifth situation also includes two sub-situations.
6. Finally, the remaining peaks not used in the above five situations will be considered as the sole signature ions.

All the peptide masses in the various situations are calculated by the weighted-average mass of the detected peaks. Table S1 summarizes the strategies that MS Annika [6], XlinkX, Maxlynx [7], MeroX RISE mode, and ECL-PF adopt based on Figure S2. ECL-PF thus adopts the most comprehensive scheme to find the potential *α/β* peptide mass region.

**Table S1:**
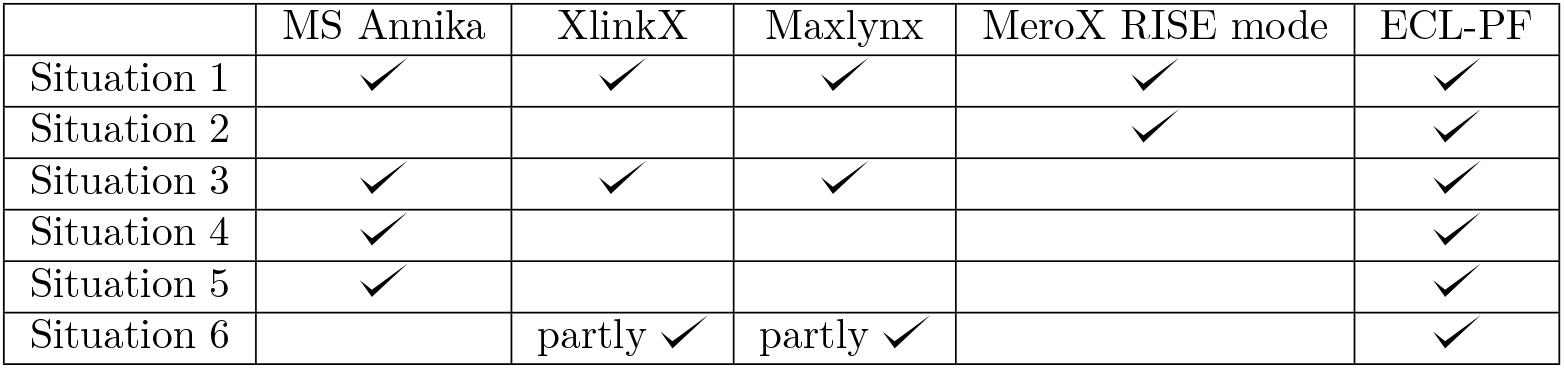
*α/β* peptide mass detection scheme that different software programs use based on the Figure S2. ‘partly’ in the table means they only use top *N* peaks instead of all the remaining peaks to derive the peptide candidates.

**Figure S2:**
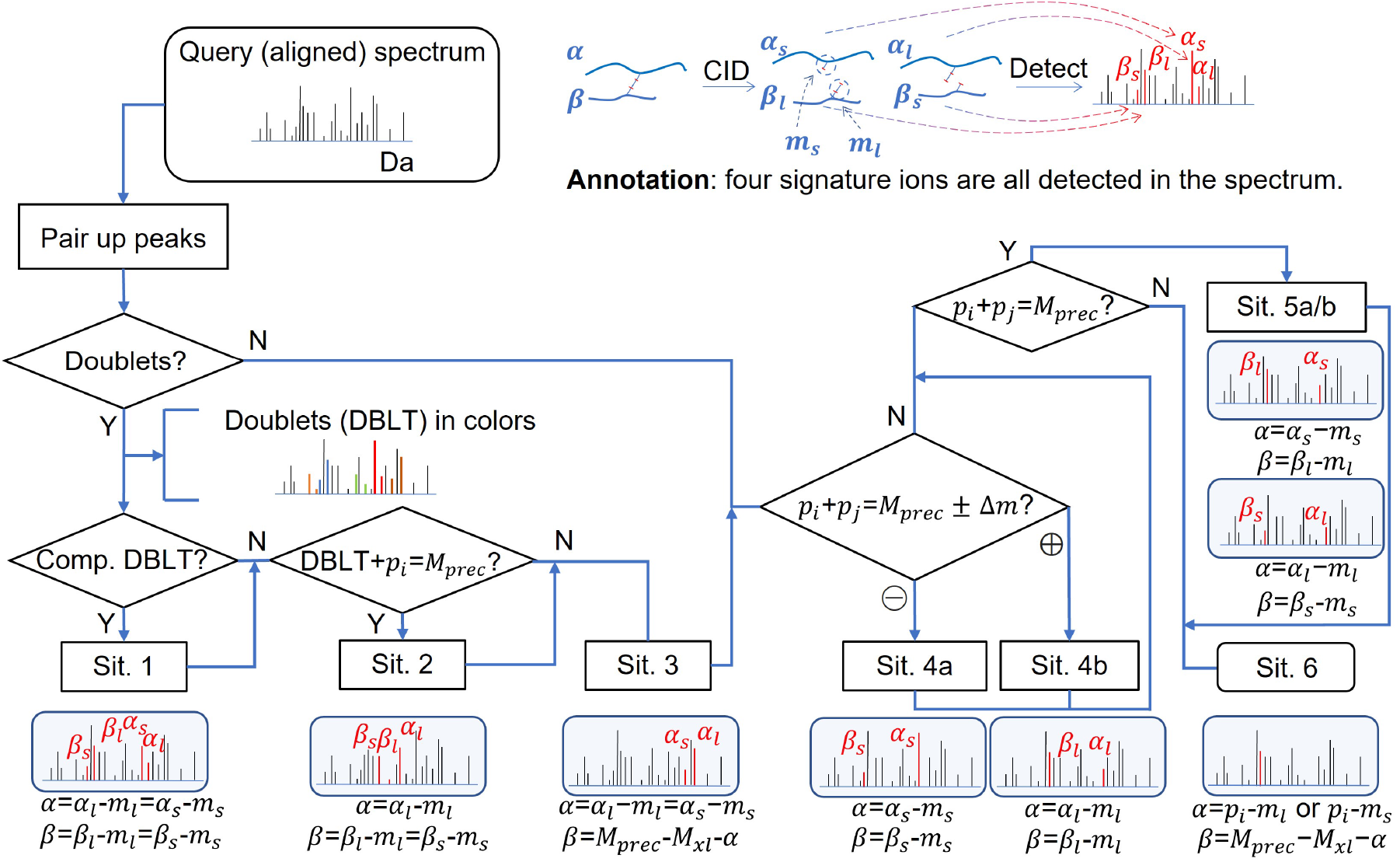
Flowchart of *α/β* peptide mass region detection. Peaks in the deisotoped spectrum (Dalton as horizontal axis) are firstly paired up to form doublets. If the distance between two peaks is equal to specific values, they are treated as a doublet (DBLT). The doublets are denoted in different colors in the figure. If there are more than two detected DBLTs, we compare any two to see if the summation of the peptide mass plus the cross-linker mass is equal to the precursor mass. If so, they are complementary doublets and this represents the appearance of signature ions of situation one (Sit. 1). The corresponding equations calculate the peptide masses. The workflow then goes to the second situation. If one doublet plus one singular peak is equal to the precursor mass, then these three peaks satisfy situation two. The doublets that cannot be used in the above two situations are singly used to derive the *α/β* peptide mass. After using all the DBLTs, we discard them in the spectrum and focus on the remaining peaks. If the summation of any two peaks is equal to the precursor mass plus/minus the delta mass, we can find the corresponding sub-situations to derive the peptide mass. If the summation of any two peaks is equal to the precursor mass alone, they satisfy situation five. Note that situation five also has two sub-situations. Last but not least, any remaining peaks that have not been used for the above five situations will be regarded as solo signature ions to infer the *α/β* peptide mass. The *α* and *β* signs in the figure stand for the *α* and *β* peptide mass. *α*_*s*_, *α*_*l*_, *β*_*s*_, and *β*_*l*_ represent an *α* peptide with a shorter residual mass, *α* peptide with a longer residual mass, *β* peptide with a shorter residual mass, and *β* peptide with a longer residual mass, respectively. *m*_*s*_ and *m*_*l*_ mean the shorter residual mass and longer residual mass of the cross-linker. Δ*m* = *m*_*l*_ − *m*_*s*_. *M*_*prec*_ and *M*_*xl*_ represent the precursor mass and cross-linker mass. *p*_*i,j*_ is any peak in the spectrum. “Comp. DBLT” is the abbreviation of complementary doublets, and Sit. 1 is situation one and so on.

ECL-PF provides an option to align the nearby spectra to include the correct peptide candidates if there does not exist a signature ion in the current spectrum. It is observed from the synthetic dataset (PXD014337) [8] that even adopting the above *α/β* peptide mass discovery scheme, some spectra still fail to be identified. The reason is that we can not detect any signature ion in the MS2 spectrum. It violates our assumption that at least one signature ion should appear in a spectrum. ECL-PF provides an option to align the nearby spectra in the *α/β* peptide mass detection process so as to use these spectra. The algorithm will not only search peaks in the current spectrum to target the *α/β* peptide mass, but also search peaks of the nearby ± 500 MS2 scans whose precursor mass is exactly the same as the current MS2 spectrum.

Though we add more peaks to guarantee our assumption, we don’t use these extra peaks in the identification process. This means that we still utilize the peaks in the current spectrum to match the peptide sequence. Table S2 shows the performance difference when we enable/disable the local alignment function in ECL-PF. In general, we can have ∼ 10% improvement in terms of the number of CSMs, and the calculated FDR barely changes.

**Table S2:**
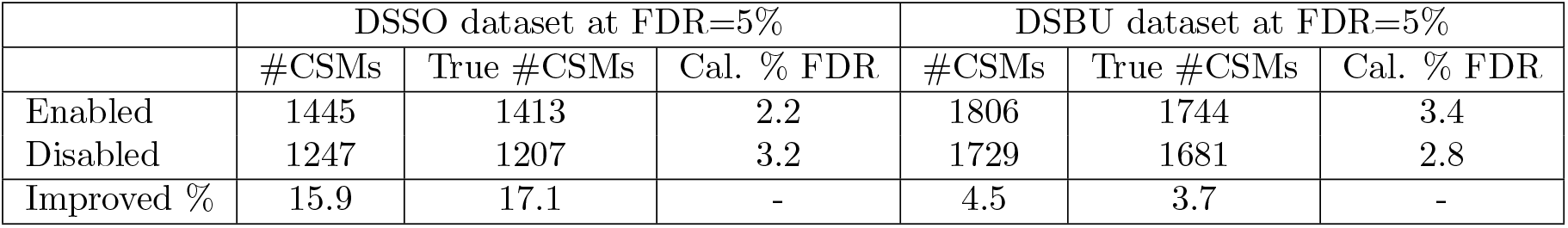
ECL-PF performance on synthetic dataset when we enable/disable local alignment function.

|  | DSSO dataset at FDR=5% |  |  | DSBU dataset at FDR=5% |  |  |
| --- | --- | --- | --- | --- | --- | --- |
|  | #CSMs | True #CSMs | Cal. % FDR | #CSMs | True #CSMs | Cal. % FDR |
| Enabled | 1445 | 1413 | 2.2 | 1806 | 1744 | 3.4 |
| Disabled | 1247 | 1207 | 3.2 | 1729 | 1681 | 2.8 |
| Improved % | 15.9 | 17.1 | - | 4.5 | 3.7 | - |

## 3 Nuts and bolts of the protein feedback pipeline

### Motivation

We observe from different datasets that, among the valid CSMs, the number of proteins corresponding to the identified peptides is very limited because not every protein can interact with others, nor does every protein have multiple available link sites inside itself. For example, in the Arabidopsis thaliana dataset [9], on average, one protein can have 6.02 different peptide sequences. But, on average, one protein can only have 1.39 unique peptide sequences. Given this information, we consider a situation in which the scoring function cannot distinguish several peptide candidates due to the lack of fragmented ions. Suppose these peptide candidates all score in a very close band. If one of the peptide candidates has many “brother” peptides that appeared in other spectra, we should regard this peptide candidate as the match for the current spectrum even though it may not have the highest score. This is the motivation for us to design a protein feedback pipeline to assist in re-matching peptide sequences with more confident ones. In Figure S3, we use an example to show how protein feedback changed the peptide ID.

**Figure S3:**
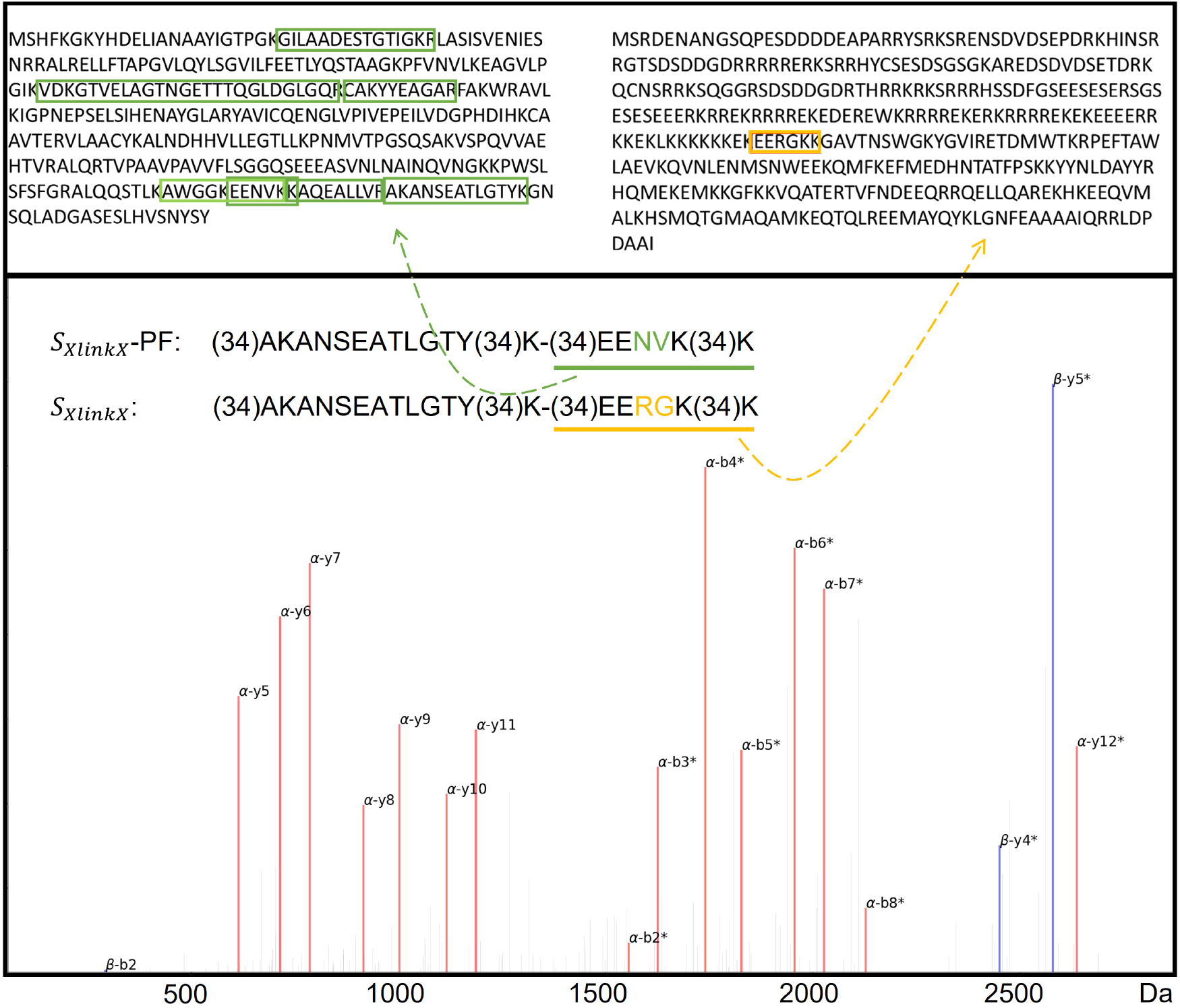
Example of the influence of protein feedback. We use XlinkX scoring function *S*_*XlinkX*_ to match the spectrum with the peptide sequence. Red peaks denote *α* matches and blue peaks denote *β* matches. Enabling and disabling protein feedback function, our program can interpret the *β* peptide differently though they both identified the same *α* peptide. We extract the corresponding proteins for these *β* peptides to find that the protein derived from *S*_*XlinkX*_ with protein feedback has many other peptides identified. But the other protein without using protein feedback is only identified once by the current spectrum.

### Choice of top hits

During the spectrum matching process, including several top hits into the peptide sequence match instead of only the top one is based on our understanding that no scoring function can work perfectly under all situations. The question is how many top hits we should include.

One solution is to use another feature to filter the top hits. After several trials, we adopt the score proportion. The score proportion is defined as the ratio of the current peptide score to the highest peptide score among the candidates (Eq. S3):

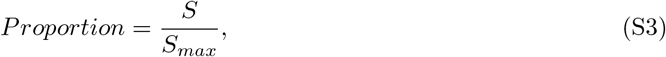

where *S* is the current peptide score and *S*_*max*_ is the highest peptide score among the current candidates.

The score proportion is similar to the delta score widely used in many software programs. The difference is that the delta score is a fixed number, while the score proportion is a dynamic criterion in which different spectra can have a different range.

To optimize the top *p* choice and proportion restriction, we use different real datasets [9–12] to run ECL-PF with different combinations of *p* and proportion. During the optimization process, we assume the higher the number of CSMs, the better the parameters. Table S3 shows the fine-tuned parameters for three scoring functions, and Figure S4 shows the results of these three scoring functions on the three datasets.

**Table S3:**
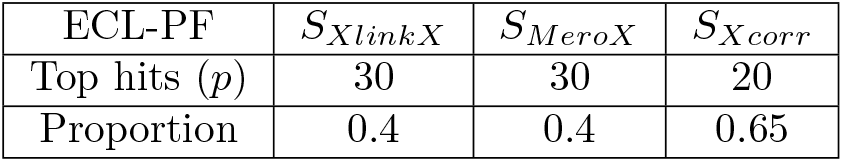
Fine-tuned parameters for different scoring functions

### Histogram of top hits and indicator function

In the middle step of the protein feedback pipeline, we retrieve all the top hits of each spectrum and plot the score distribution. This step provides global information. Figure S5 shows the histograms of different scoring functions. The target results and decoy results [13] are plotted in red and blue colors, respectively. It is obvious that true (target) results tend to have higher scores than the false (decoy) results, regardless of what scoring function we are using. Thus, we can use this consistent property to re-select the peptide sequence for different scoring functions.

To determine the significant domain in the histogram, we employ an indicator function. From Figure S5, we can observe that the position at twice the median number can roughly filter out all the decoy results. Alternatively, the 98th percentile is also a good cutoff choice. Please note that ECL-PF discards all the results with the 0 score when calculating the threshold.

### Protein feedback core: Counting function

One important purpose of designing the protein feedback pipeline is to apply it to all existing scoring functions. Two problems will arise if we directly accumulate each peptide’s score onto that corresponding protein.

- If the score is not designed linearly proportionately to the peptide confidence, the protein score will be biased.
- Protein score will not be comparable among the different scoring functions.

Therefore, we need to design the counting method based on the following rules:

1. Two peptides identified in one protein should count as more than one peptide identified twice in one protein.
2. If one peptide is identified several times in one protein, the count should increase but not become larger than some threshold.
3. If several peptides are identified in one spectrum that is all ranked in the 1st position, they should all be counted into the protein score but have a uniform weight of 1*/n*, where *n* is the number of peptides ranked in the 1st position.
4. If no peptide is identified in a protein, the protein score should be 0.

**Figure S4:**
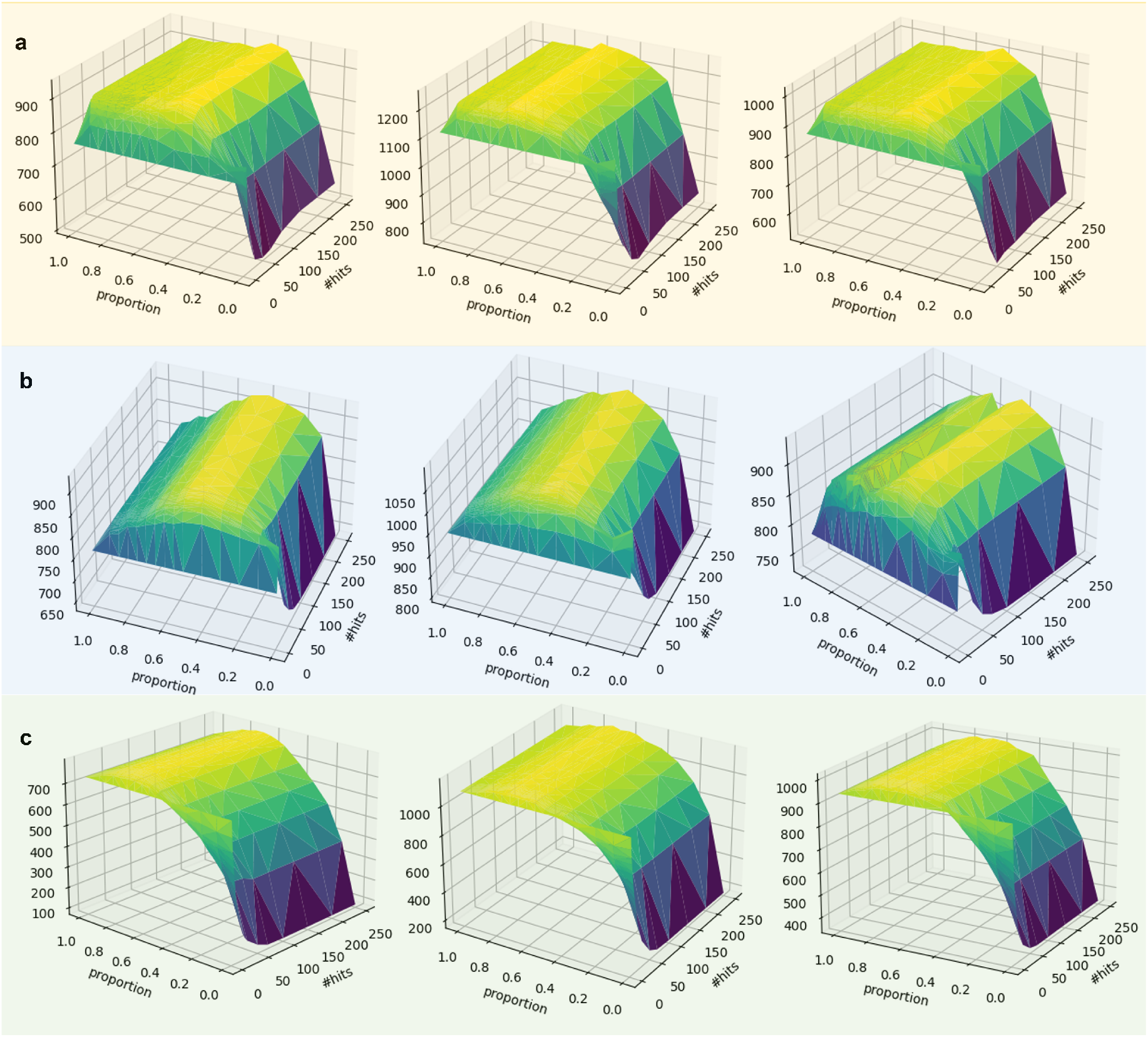
#CSM as a function of #top hits and score proportion for three different software with three different datasets. The surface is almost flat when the number of top hits goes larger and larger for every scoring functions. In order to save cache storage, we choose the least number of top hits at the plateau. (a) XlinkX scoring function result. The plateau is roughly located at the *p* = 30 and proportion = 0.4. (b) MeroX scoring function result. The plateau is roughly located at the *p* = 30 and proportion = 0.4. (c) Xcorr scoring function result. The plateau is roughly located at the *p* = 20 and proportion = 0.65.

**Figure S5:**
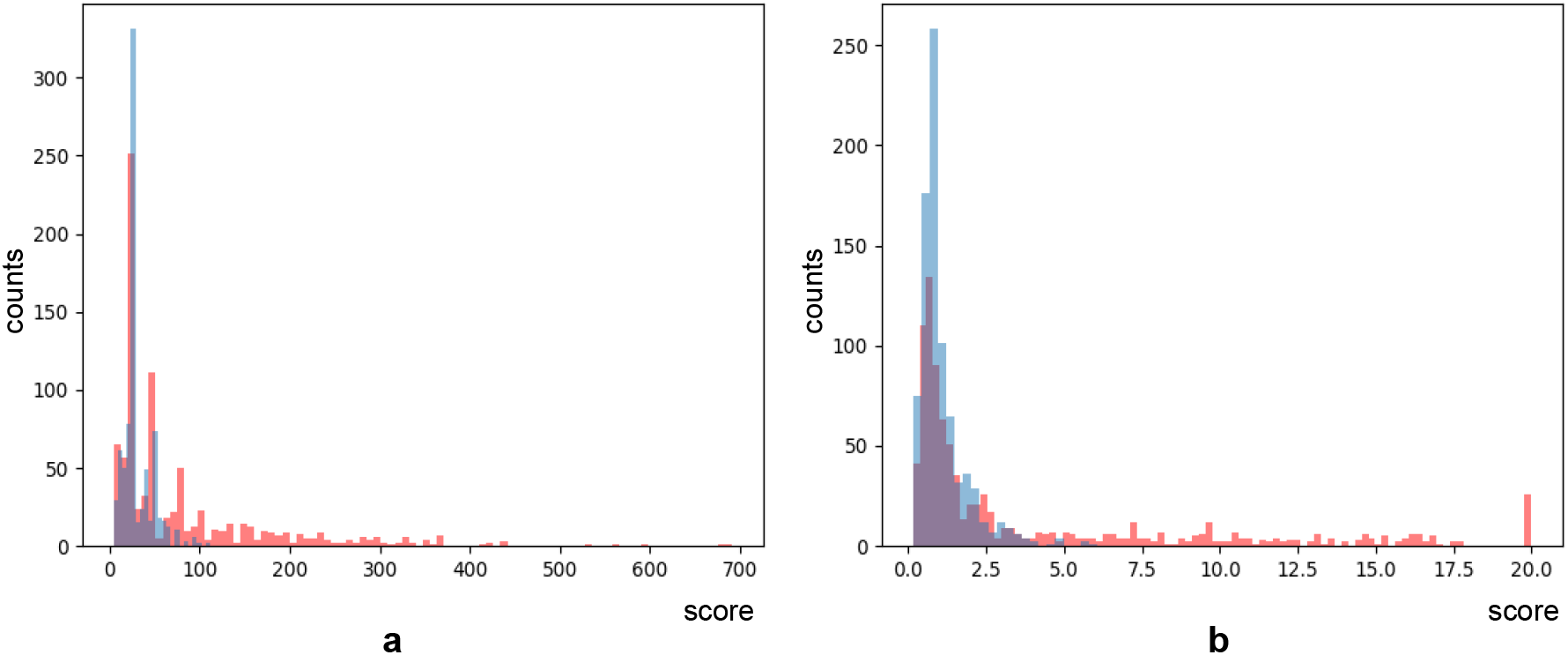
Score histogram of top hits. Target results (red) are drawn separately from the decoy results (blue). The two sub-figures show the histograms for the same dataset. (a) Score histogram of Xcorr scoring function. (b) Score histogram of XlinkX scoring function. The results show that a properly designed scoring function always tends to match the true peptides with higher scores. This provides us with a common standard to select significant peptides.

Based on the above rules, we construct a counting function *C*(*x*) using a skewed form of the hyperbolic tangent function and define the final protein score PS as the sum of counting functions:

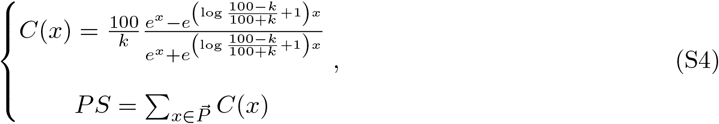

where *k* is the number of link sites in the individual protein and 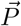 is the protein vector. Figure S6 shows how it looks with three different k settings (k=20, k=25, k=50).

This counting function has properties that satisfy all of our rules.

- *C*(*x*) + *C*(*y*) ≥ *C*(*x* + *y*) for any *x, y* ≥ 0. This property satisfies rule 1.
- *C*(*x*) has an upper bound of 100*/k* no matter how many times the same peptides are identified. Here, *k* is the total number of link sites in a protein. This guarantees that for different proteins in the database, each protein score is comparable to others, which satisfies rule 2.
- The counting function is a continuous function instead of a discrete function, which satisfies rule 3.
- The counting function always passes through (0,0), which means that if no peptides are identified in one protein, the protein score should be 0. This satisfies rule 4. The function also passes through (1,1) all the time. If one peptide is identified only once, then the output should be exactly 1.

**Figure S6:**
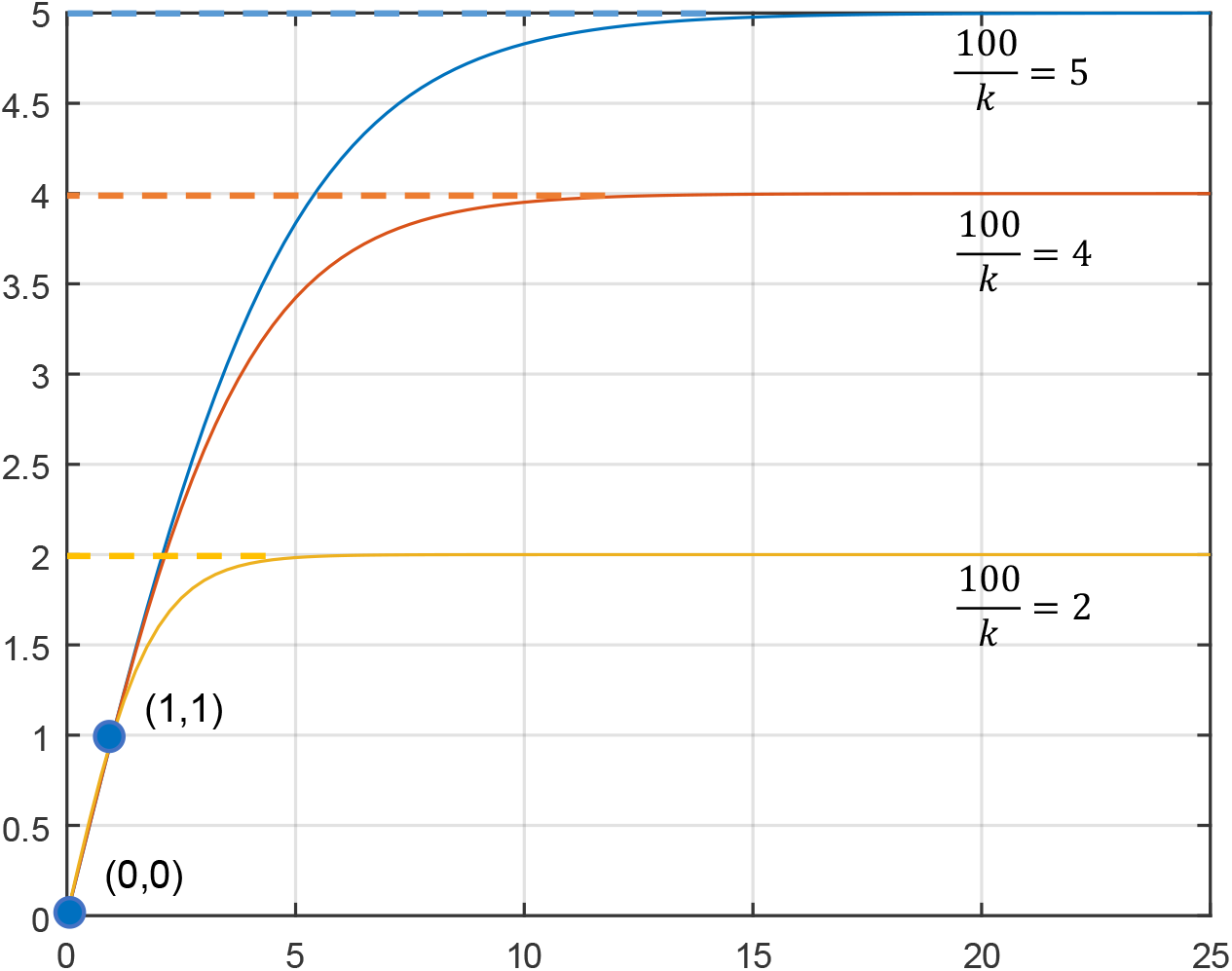
Visualization of counting function when *k* = 20, 25, and 50. The function always passes through (0,0) and (1,1) and has the upper bound of 100*/k*.

### Protein feedback on different scoring functions

XlinkX [2, 14] and MeroX [15] are two widely used software in cleavable XL-MS. SEQUEST’s scoring function [16], Xcorr, is also a simple and robust scoring function. We integrated these three scoring functions into our ECL-PF. By disabling and enabling the protein feedback module, we observe that the performance of all these scoring functions improves. On average, the protein feedback pipeline can provide 29.7% more CSMs, shown in Table S4. The first column of the table also shows the benchmark results of MeroX and MaxLynx.

**Table S4:** Generalizability and effectiveness of protein feedback on different scoring functions

| Benchmark | ECL performance |  |  |
| --- | --- | --- | --- |
| MeroX | ECL(XlinkX) | ECL(MeroX) | ECL(Xcorr) |
| 28826 | 35990 | 28391 | 35031 |
| MaxLynx | ECL-PF(XlinkX) | ECL-PF(MeroX) | ECL-PF(Xcorr) |
| 31754 | 48075 | 37329 | 43292 |
| Improved % | 33.59% | 31.48% | 23.58% |

## 4 Precursor mass refinement

Precursor mass refinement is prevalent in XL-MS software, and correct precursor mass information is key to the success of peptide identification. ECL-PF approximates the theoretical isotope distribution [17] and compares it to the experimental MS1 spectrum to derive the correct precursor mass. Figure S7 shows the workflow of this module. The nearby MS1 spectra of the current MS2 spectrum are extracted and merged together to form a new spectrum. We use four preceding MS1 and three subsequent MS1 to implement this step and also restrict the scan event. A fused isotope cluster is more similar to the theoretical isotope cluster than the original result, shown in Figure S7(b). After obtaining the new spectrum, we compare the theoretical isotope cluster with each possible position in the new MS1 and calculate the corresponding Pearson correlation coefficient using Eq. S5. The best fit is our new calculated precursor mass result. In ECL-PF, if we cannot observe any coefficient larger than the defined threshold (0.9 by default), we then retain the originally provided precursor mass from the RAW file.

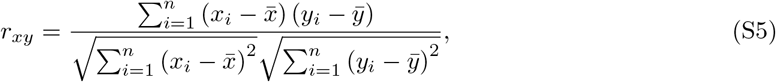

where *x* and *y* stand for the theoretical and experimental isotope clusters, respectively.

Using different datasets, we observed that precursor mass refinement module can improve the number of CSMs by ∼ 30% in Figure S8. Table S5 shows even higher improvements using the synthetic dataset (PXD014337).

**Table S5:** ECL-PF performance on synthetic dataset when we enable/disable precursor mass refinement function.

|  | DSSO dataset at FDR=5% |  |  | DSBU dataset at FDR=5% |  |  |
| --- | --- | --- | --- | --- | --- | --- |
|  | #CSMs | True #CSMs | Cal. % FDR | #CSMs | True #CSMs | Cal. % FDR |
| Enabled | 1445 | 1413 | 2.2 | 1806 | 1744 | 3.4 |
| Disabled | 826 | 763 | 7.6 | 1500 | 1450 | 3.3 |
| Improved % | 74.9 | 85.2 | - | 20.4 | 20.2 | - |

*α/β* mass detection scheme, protein feedback, and precursor mass refinement module are three major functions that improve the performance (sensitivity) of ECL-PF. Figure S8 summarizes how much each module contributes to the final improvements. On average, the result shows that the number of CSMs improves two-fold after adopting these modules.

**Figure S7:**
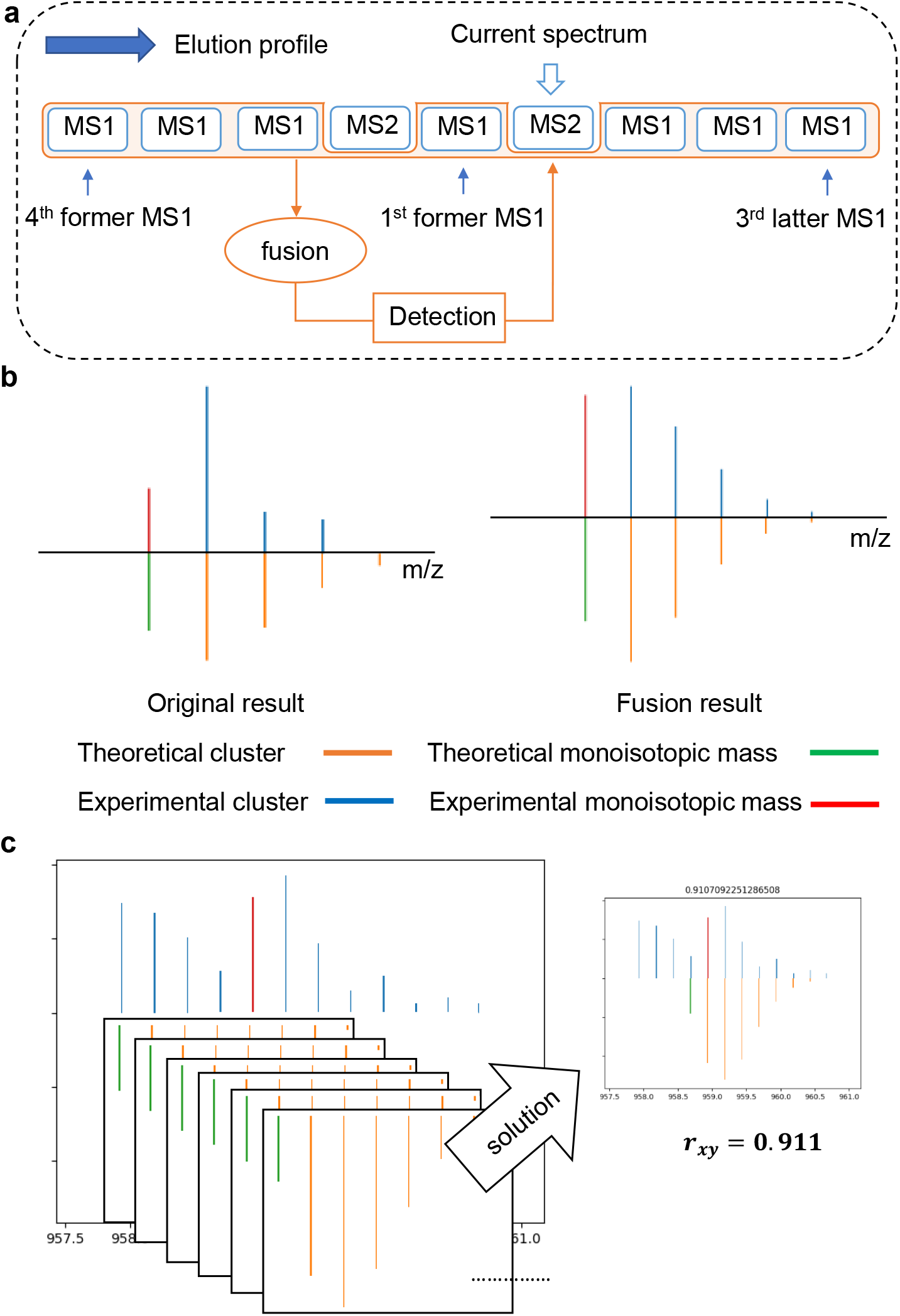
Workflow of precursor mass refinement module in ECL-PF. (a) Nearby MS1 spectra are merged to generate a complete isotope cluster. (b) Comparison of isotope cluster between single (original) MS1 and merged MS1. The fusion result has a better shape and is more similar to the theoretical isotope cluster. (c) Theoretical isotope cluster is compared with each possible position in MS1 and Pearson coefficient is used to measure the similarity.

**Figure S8:**
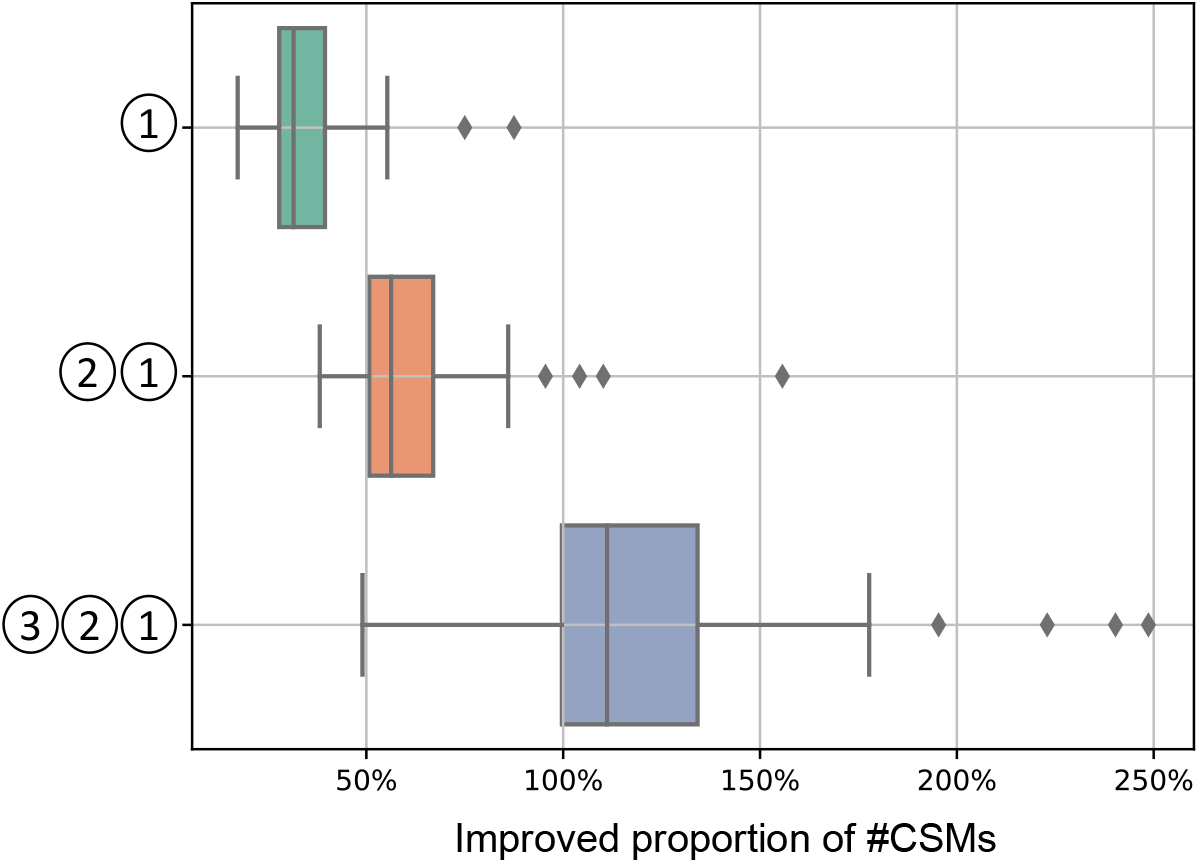
Performance improvements using different modules in ECL-PF. Precursor mass refinement module ①, exhaustive *α/β* mass detection module ② and protein feedback module ③ are three major functions that improve the sensitivity of cross-linking identifications. The box plot shows how much each part contributes to the overall improvement of sensitivity. We use *S*_*XlinkX*_ score function with precursor mass provided by the data to match the peptide sequences, and only consider situation 1 and situation 3 to detect *α/β* peptide. We then treat this result as the baseline. On average, after adopting ①, the number of CSMs increases by 34.4%. Further integrating ②, the number of CSMs increases by 61.1%. Finally, combining ③, the number of CSMs moves up to 120.7%.

## 5 Hybrid simulated dataset

Based on the properties of XL-MS, we generate a hybrid simulated dataset that is close to a real experimental one. Figure S9 shows the procedure.

For any real dataset, we use several software programs to analyze it simultaneously and treat the overlapping identified spectra as the source data. For each spectrum in the source data, we categorize the peaks into signal and noise. For signal peaks, a/b/c ions, x/y/z ions, internal ions and neutral loss ions could possibly be included, depending on the data type. For noise peaks, all other peaks excluding the signal, doublets, reporter ions and precursor ions could be included. The signals can be further divided into *α* peptide peaks and *β* peptide peaks. After extracting signal and noise, we combine every *α* signal from one spectrum with every *β* signal from another spectrum plus their noise to produce a new spectrum. In this way, we can augment *n* spectra with ground truth to *n*^2^ spectra. The advantage of this method is that we don’t need to generate the dataset from scratch.

In this paper, the source data are derived from ECL-PF and MeroX results. b/y ions are considered as the signal only. The noise proportion is tuned for different simulated subgroups, as shown in Table S6. For example, when the noise proportion is 40%, we randomly pick 40% of noise peaks from the original source to generate the new spectrum.

**Figure S9:**
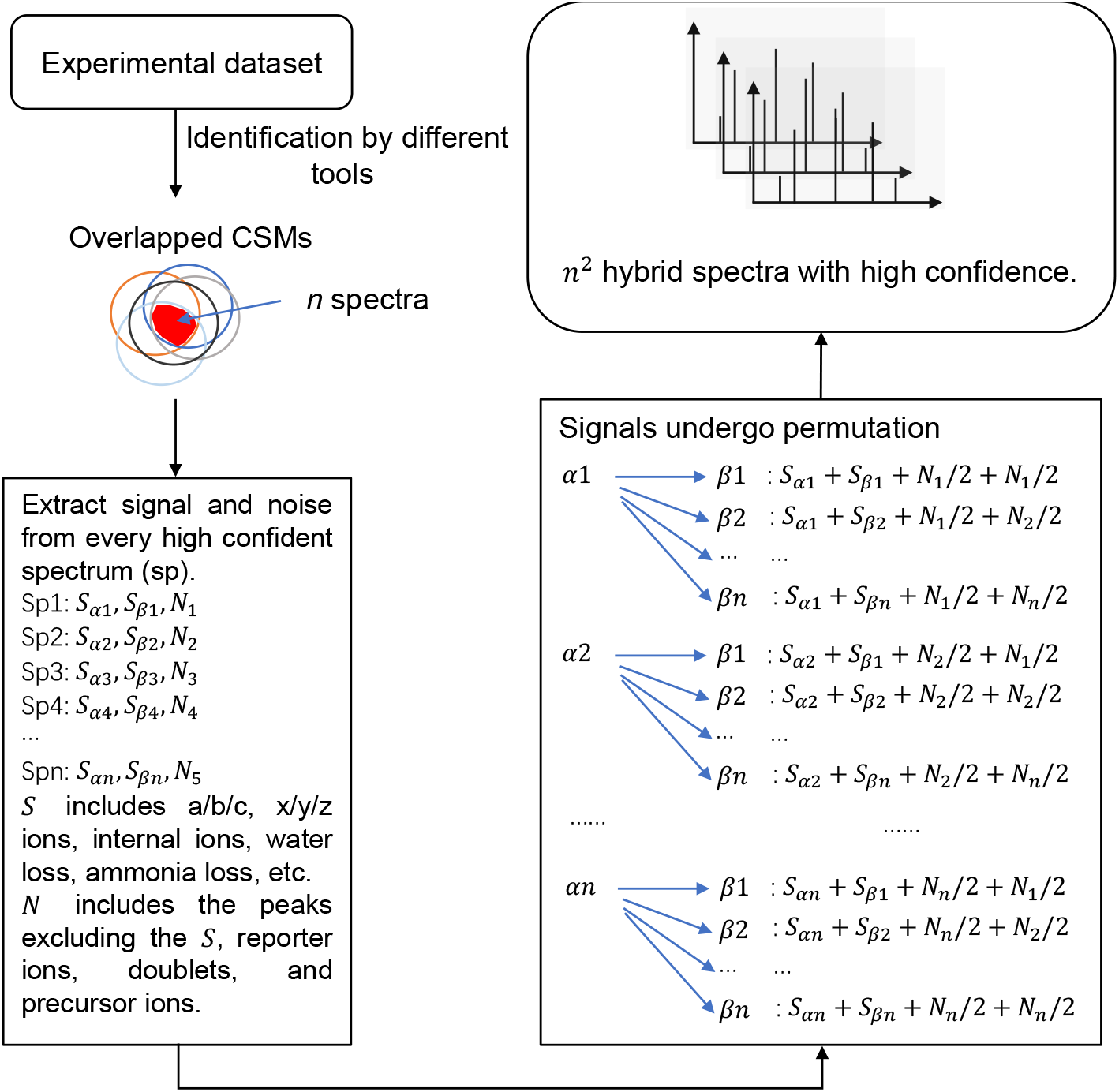
Procedure of hybrid dataset simulation. Experimental datasets are analyzed by several software programs and the overlapping spectra are regarded as the source data. Suppose there are *n* spectra from the source. For every individual spectrum from the source, we separate the signal from noise. The types of the signal may include a/b/c ions, x/y/z ions, internal ions and neutral loss ions. Noise peaks are the other peaks, excluding signal, doublets, reporter ions and precursor ions. The source ions then undergo permutations to generate *n*^2^ new spectra. The noise proportion in the middle step can be tuned.

**Table S6:** Hybrid simulated dataset composition in this paper

| Source | #Spectra | Noise proportion | Signal proportion |
| --- | --- | --- | --- |
| DSBU dataset | 14161 | 0% | 100% |
|  | 13924 | 20% | 100% |
|  | 13924 | 40% | 100% |
|  | 13924 | 60% | 100% |
|  | 13924 | 80% | 100% |
|  | 14884 | 100% | 100% |
| DSSO dataset | 4225 | 0% | 100% |
|  | 4225 | 20% | 100% |
|  | 4225 | 40% | 100% |
|  | 4225 | 60% | 100% |
|  | 4225 | 80% | 100% |
|  | 4096 | 100% | 100% |

## 6 Data descriptions, parameter settings and supporting figures

Table S7 shows the concrete information for all the datasets used in this paper: PXD014337 [8], PXD011861 [18], PXD016963 [19], PXD020014 [20] and PXD012546 [10].

**Table S7:**
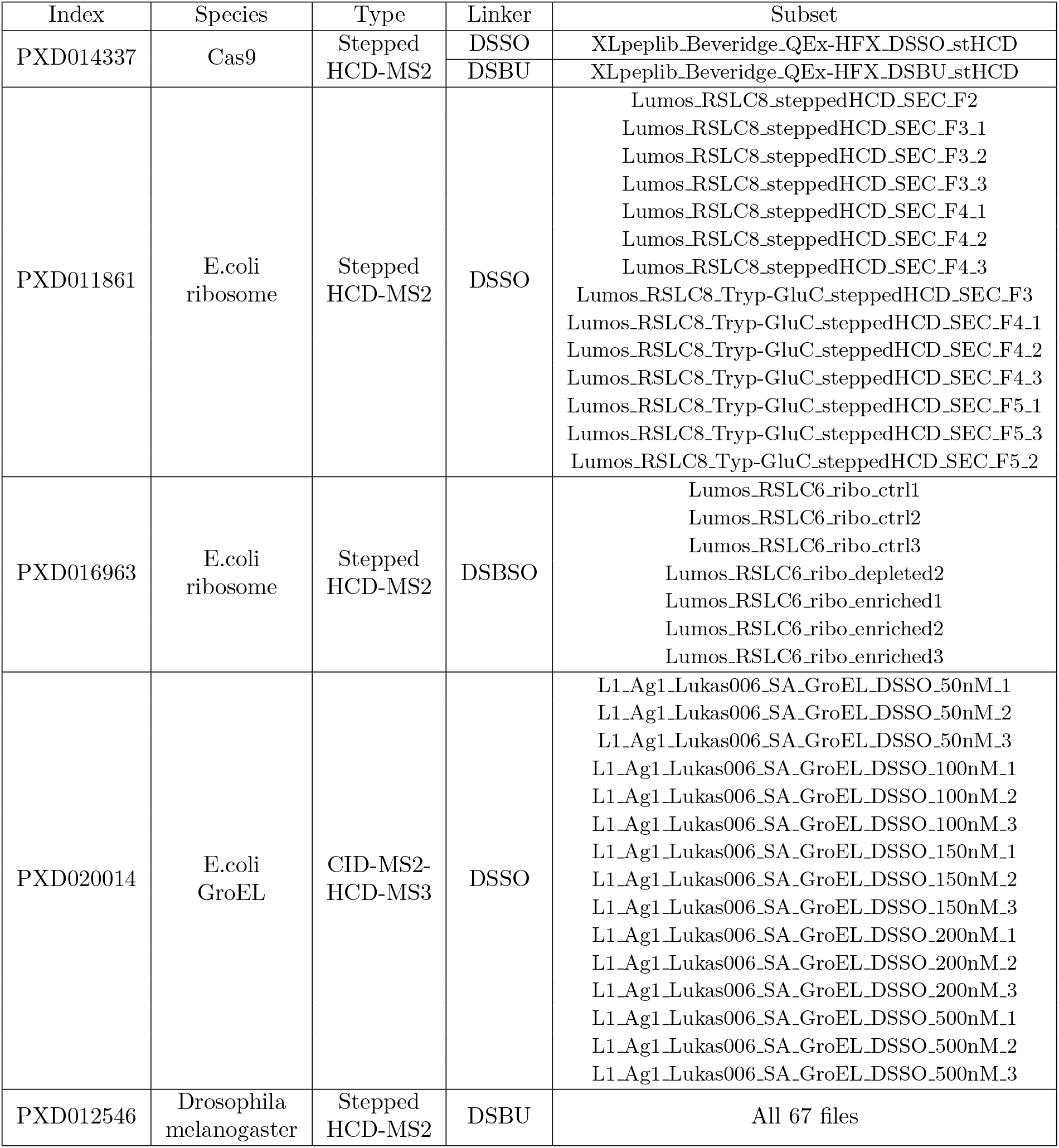
Datasets used in this paper. Though some datasets have MS3 spectra, no software will use them in the identification process for fair comparison.

**Table S8:** Parameters in the PXD014337 synthetic dataset

| PXD014337 | ECL-PF | MaxLynx | MeroX | MS Annika | XlinkX |
| --- | --- | --- | --- | --- | --- |
| Enzyme | Trypsin | Trypsin | Trypsin | Trypsin | Trypsin |
| Miss_cleavages | 2 | 2 | 2 | 2 | 2 |
| Min_length | 5 | 5 | 5 | 5 | 5 |
| Modifications | C+57.02Da<br>(fixed)<br>M+15.99Da<br>(variable) | C+57.02Da<br>(fixed)<br>M+15.99Da<br>(variable) | C+57.02Da<br>(fixed)<br>M+15.99Da<br>(variable) | C+57.02Da<br>(fixed)<br>M+15.99Da<br>(variable) | C+57.02Da<br>(fixed)<br>M+15.99Da<br>(variable) |
| MS1 tolerance | 5ppm | 5ppm | 5ppm | 5ppm | 5ppm |
| MS2 tolerance | 20ppm | 20ppm | 20ppm | 20ppm | 20ppm |
| Linker info | DSSO $m_{xl} = 158.0038, m_s = 54.0106, m_l = 85.9826$<br>DSBU $m_{xl} = 196.0848, m_s = 85.0528, m_l = 111.032$ | | | | |
| Link site | K | K | K | K | K |

**Table S9:** Parameters in the hybrid simulated dataset

| PXD014337 | ECL-PF | MeroX |
| --- | --- | --- |
| Enzyme | Trypsin | Trypsin |
| Miss_cleavages | 3 | 3 |
| Min_length | 5 | 5 |
| Modifications | C+57.02Da<br>(fixed)<br>M+15.99Da<br>(variable) | C+57.02Da<br>(fixed)<br>M+15.99Da<br>(variable) |
| MS1 tolerance | 5ppm | 5ppm |
| MS2 tolerance | 20ppm | 20ppm |
| Linker info | DSSO $m_{xl} = 158.0038, m_s = 54.0106, m_l = 85.9826$<br>DSBU $m_{xl} = 196.0848, m_s = 85.0528, m_l = 111.032$ | |
| Link site | K | K |

**Table S10:** Parameters in the PXD011861 synthetic dataset

| PXD011861 | ECL-PF | MaxLynx | MeroX |
| --- | --- | --- | --- |
| Enzyme | Trypsin | Trypsin | Trypsin |
| Miss_cleavages | 4 | 4 | 4 |
| Min_length | 5 | 5 | 5 |
| Modifications | C+57.02Da<br>(fixed)<br>M+15.99Da<br>(variable) | C+57.02Da<br>(fixed)<br>M+15.99Da<br>(variable) | C+57.02Da<br>(fixed)<br>M+15.99Da<br>(variable) |
| MS1 tolerance | 10ppm | 10ppm | 10ppm |
| MS2 tolerance | 20ppm | 20ppm | 20ppm |
| Linker info | DSSO $m_{xl} = 158.0038, m_s = 54.0106, m_l = 85.9826$ | | |
| Link site | N-term, K | N-term, K | N-term, K |

**Table S11:**
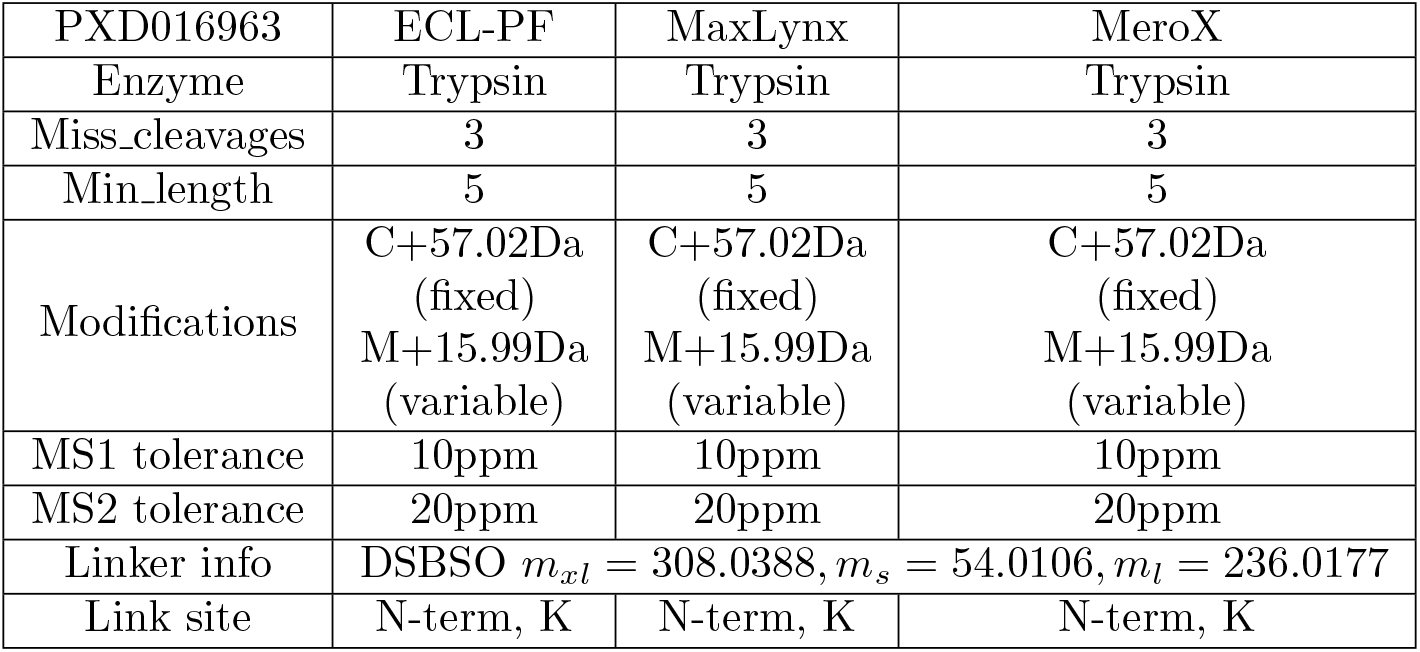
Parameters in the PXD016963 synthetic dataset

| PXD016963 | ECL-PF | MaxLynx | MeroX |
| --- | --- | --- | --- |
| Enzyme | Trypsin | Trypsin | Trypsin |
| Miss_cleavages | 3 | 3 | 3 |
| Min_length | 5 | 5 | 5 |
| Modifications | C+57.02Da<br>(fixed)<br>M+15.99Da<br>(variable) | C+57.02Da<br>(fixed)<br>M+15.99Da<br>(variable) | C+57.02Da<br>(fixed)<br>M+15.99Da<br>(variable) |
| MS1 tolerance | 10ppm | 10ppm | 10ppm |
| MS2 tolerance | 20ppm | 20ppm | 20ppm |
| Linker info | DSBSO $m_{xl} = 308.0388, m_s = 54.0106, m_l = 236.0177$ | | |
| Link site | N-term, K | N-term, K | N-term, K |

**Table S12:** Parameters in the PXD020014 synthetic dataset

| PXD020014 | ECL-PF | MaxLynx | MeroX |
| --- | --- | --- | --- |
| Enzyme | Trypsin | Trypsin | Trypsin |
| Miss_cleavages | 2 | 2 | 2 |
| Min_length | 5 | 5 | 5 |
| Modifications | C+57.02Da<br>(fixed)<br>M+15.99Da<br>(variable) | C+57.02Da<br>(fixed)<br>M+15.99Da<br>(variable) | C+57.02Da<br>(fixed)<br>M+15.99Da<br>(variable) |
| MS1 tolerance | 10ppm | 10ppm | 10ppm |
| MS2 tolerance | 20ppm | 20ppm | 20ppm |
| Linker info | DSSO $m_{xl} = 158.0038, m_s = 54.0106, m_l = 85.9826$ | | |
| Link site | N-term, K | N-term, K | N-term, K |

**Table S13:** Parameters in the PXD012546 synthetic dataset

| PXD012546 | ECL-PF | MaxLynx | MeroX |
| --- | --- | --- | --- |
| Enzyme | Trypsin | Trypsin | Trypsin |
| Miss_cleavages | 3 | 3 | 3 |
| Min_length | 5 | 5 | 5 |
| Modifications | C+57.02Da<br>(fixed)<br>M+15.99Da<br>N-term+42.0367Da<br>(variable) | C+57.02Da<br>(fixed)<br>M+15.99Da<br>N-term+42.0367Da<br>(variable) | C+57.02Da<br>(fixed)<br>M+15.99Da<br>N-term+42.0367Da<br>(variable) |
| MS1 tolerance | 5ppm | 5ppm | 5ppm |
| MS2 tolerance | 15ppm | 15ppm | 15ppm |
| Linker info | DSBU $m_{xl} = 196.0848, m_s = 85.0528, m_l = 111.032$ | | |
| Link site | N-term, K | N-term, K | N-term, K |

**Figure S10:**
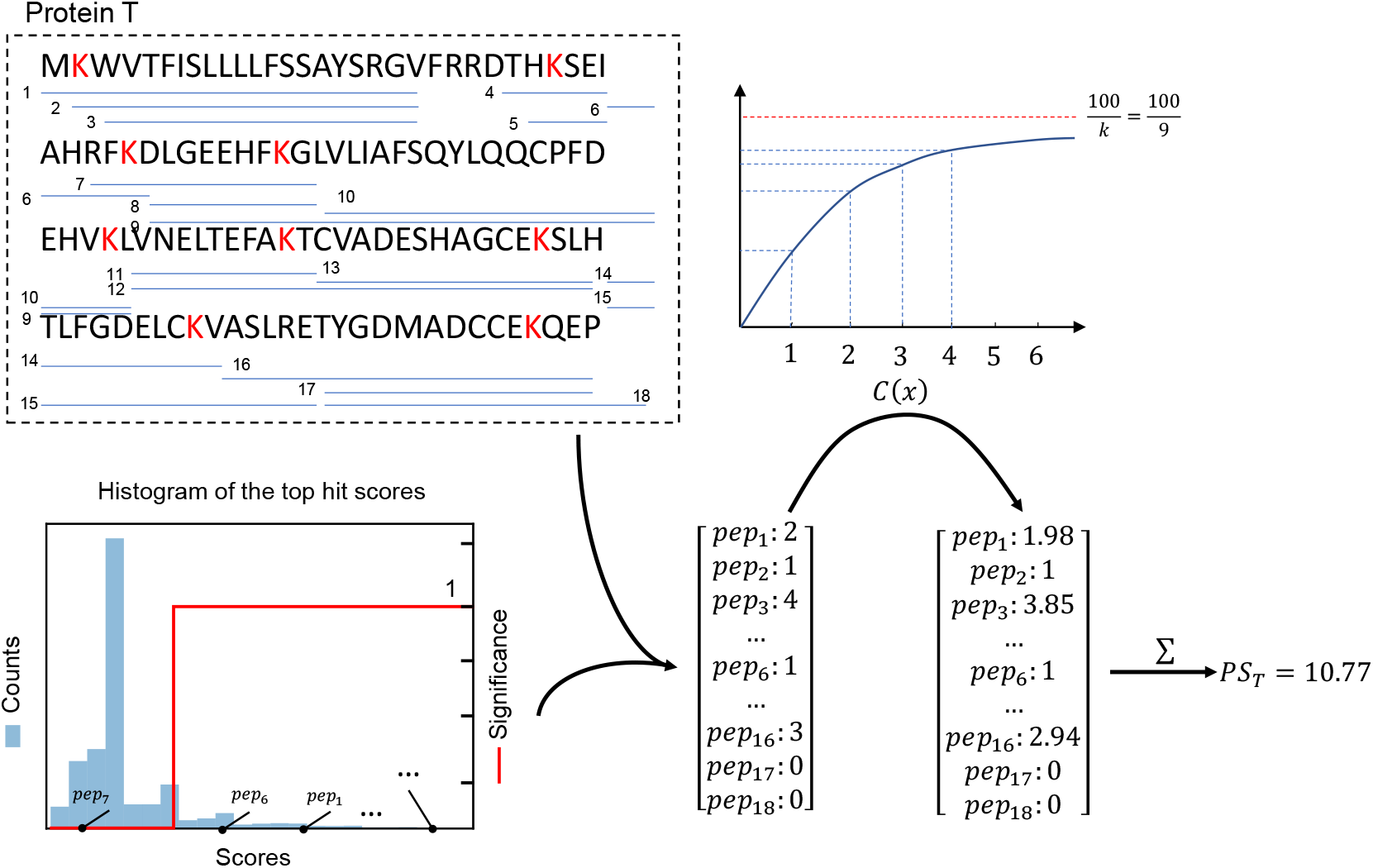
Concrete example to calculate protein score. Protein *T* can be digested into 18 distinct peptide sequences under the one missed cleavage setting. Therefore, the protein vector length is 18. From the histogram of top hit scores, only *pep*_1_, *pep*_2_, *pep*_3_, *pep*_6_, and *pep*_16_ can be found in the significant region. We initialize the protein vector with their numbers. Then adopting counting function *C*(*x*), we transfer the protein vector into a new one, where parameter *k* is 9 in the *C*(*x*). Last, we sum all the elements in the vector to calculate the final score of *T*.

---

**Figure S11:**
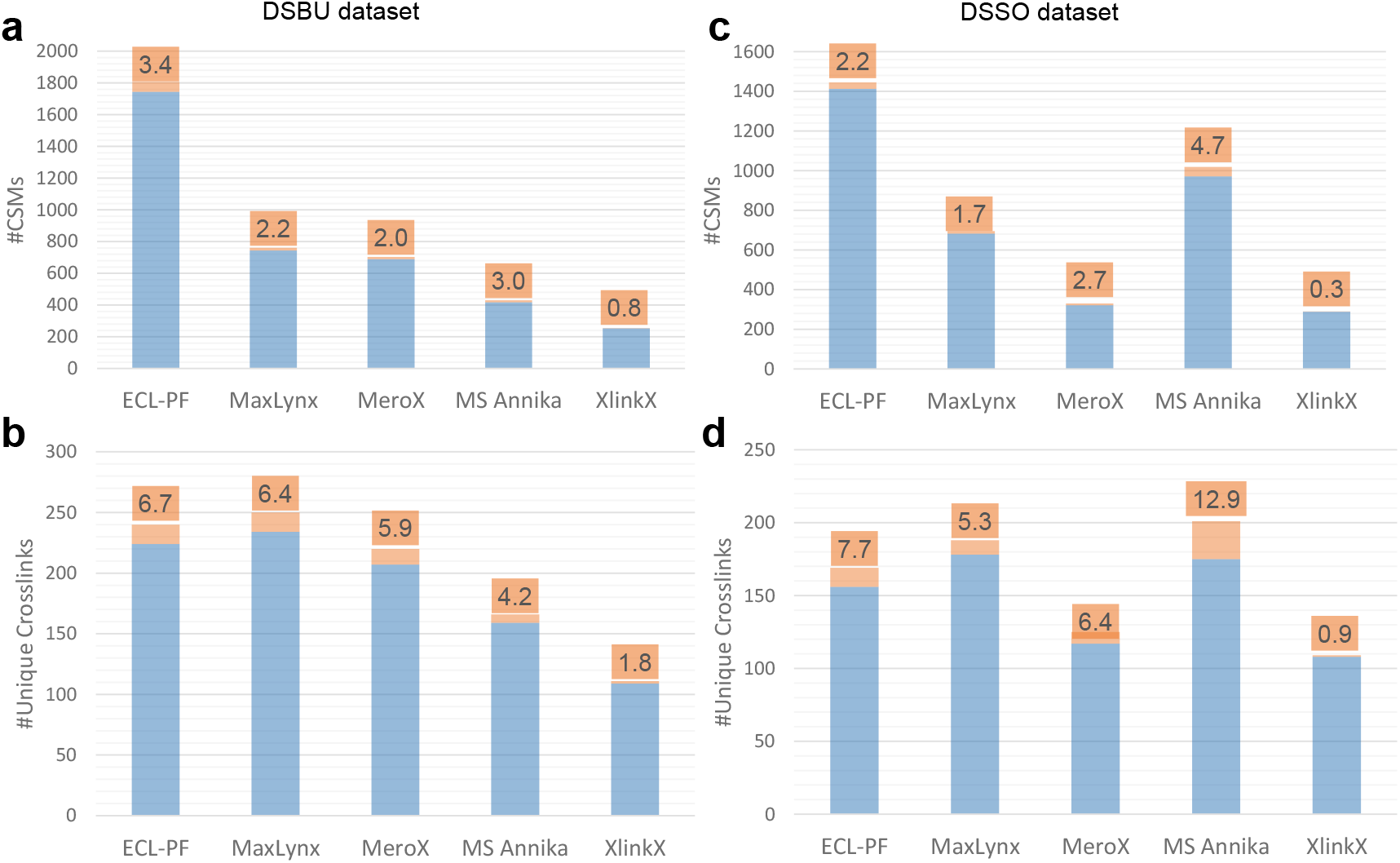
Software comparison on synthetic datasets at FDR=1%. CSMs and unique cross-link results for DSBU and DSSO datasets are plotted. True positive results are denoted by the blue bar. False positives are denoted by the orange bar, and the proportion is labeled. On average, ECL-PF identifies 2 ∼ 3 times as many CSMs as the other software. Due to the limited number of synthesized peptides in the samples, ECL-PF doesn’t outperform all the other software in terms of the unique cross-links but is in the same range as the best one.

